# Generalizable Morphological Profiling of Cells by Interpretable Unsupervised Learning

**DOI:** 10.1101/2024.09.24.614684

**Authors:** Rashmi Sreeramachandra Murthy, Shobana V. Stassen, Dickson M. D. Siu, Michelle C. K. Lo, Gwinky G. K. Yip, Kevin K. Tsia

## Abstract

The intersection of advanced microscopy and machine learning is revolutionizing cell biology into a quantitative, data-driven science. While traditional morphological profiling of cells relies on labor-intensive manual feature extraction susceptible to biases, deep learning offers promising alternatives but struggles with the interpretability of its black-box operation and dependency on extensive labeled data. We introduce MorphoGenie, an unsupervised deep-learning framework designed to address these challenges in single-cell morphological profiling. Enabling disentangled representation learning integrated with high-fidelity image reconstructions, MorphoGenie possesses a critical attribute to learn a compact, generalizable and interpretable latent space. This facilitates the extraction of biologically meaningful features without human annotation, additionally overcoming the "curse of dimensionality" inherent in manual methods. Unlike prior models, MorphoGenie introduces a systematic approach to mapping disentangled latent representations to fundamental hierarchical morphological attributes, ensuring both semantic and biological interpretability. Moreover, it adheres to the concept of combinatorial generalization—a core principle of human intelligence— which greatly enhances the model’s capacity to generalize across a broad spectrum of imaging modalities (e.g., quantitative phase imaging and fluorescence imaging) and experimental conditions (ranging from discrete cell type/state classification to continuous trajectory inference). The framework offers a new, generalized strategy for unbiased and comprehensive morphological profiling, potentially revealing insights into cellular behavior in health and disease that might be overlooked by expert visual examination.

## Introduction

Recent advances in microscopy have revolutionized cell biology by transforming it into a data-driven science. This transformation allows researchers to explore the rich structural and functional traits of cell morphology, providing valuable insights into cell health, disease mechanisms, and cellular responses to chemical and genetic perturbations. In recent years, we have witnessed remarkable growth in open image data repositories [1-5] and the development of powerful machine learning methods for analyzing cellular morphological fingerprints (or profiles). There is mounting evidence that these morphological profiles can reveal critical information about cell functions and behaviors, often remaining hidden in molecular assays. Notably, it has been shown that cell morphology and gene expression profiling provide complementary information in genetic and chemical perturbations [6-8].

Traditional morphological profiling methods rely on manual feature extraction, which can be labor-intensive, require domain expertise, and often lack scalability and generalizability across different imaging modalities. Conventionally, features are crafted based on cellular shape, size, texture, and pixel intensities to assign a unique identity to each cell. Extracting hundreds to thousands of morphological features from a single image allows the investigation of complex cellular properties with high discriminative power, such as responses to drug treatments [9, 10]. However, manual feature extraction methods are susceptible to the "curse of dimensionality" and may introduce biases, as the selected features might not fully represent the data.

Deep learning techniques that employ supervised or weakly supervised learning have shown promise in delivering more accurate image classification [11]. However, these methods require large-scale, expert labeling or annotation of training datasets, which can be time-consuming and subject to human biases [12]. Additionally, the performance of deep learning is often hindered by its lack of interpretability. An ideal cell morphology profiling strategy should generate features without overly depending on human knowledge, making inferences based solely on the images themselves, free from any a priori assumptions. Adopting such an approach has shown effective to extract subtle cellular features that are obscured through manual feature extraction; and to offer a more unbiased analysis of cellular morphology, overcoming the limitations associated with manual annotation and expert knowledge. At the same time, the deep-learned morphological profile should effectively be interpretable (or explainable) in order to improve the deep learning model transparency and gain credibility, which is particularly important in biomedical diagnosis [13, 14].

Unsupervised deep generative networks, notably variational autoencoders or VAEs [15], have gained widespread success in learning interpretable latent representations for downstream analysis and providing insights into neural network model learning. Autoencoders learn to compress input data into a lower-dimensional representation (encoding) and then reconstruct the input image data from this lower-dimensional representation (decoding), while learning the latent representations. Despite their potential, autoencoders often face limitations in lossy image reconstructions - making it difficult to assess how well the model can learn a good probabilistic latent representation of the image data. While previous works have employed VAE variants for unsupervised and self-supervised learning of cellular image datasets to reveal cellular dynamics and attempted to interpret the learned latent space [16-18], they have not established a direct and systematic mapping between the learned latent space and interpretable morphological features. This highlights the need for further research to overcome these limitations and enhance the morphological profiling of cells.

We present MorphoGenie, a new deep-learning framework for unsupervised, interpretable single-cell morphological profiling and analysis to address the abovementioned challenges. MorphoGenie distinguishes itself from previous works with three key attributes: (1) *High-fidelity Image Reconstruction*: MorphoGenie utilizes a hybrid architecture that capitalizes on the unparalleled strengths of the variant of VAEs and generative adversarial networks (GANs) to achieve interpretable, high-quality cell image generation [19]. (2) *Interpretability*: MorphoGenie learns a compact, interpretable, and transferable disentangled representation for single-cell morphological analysis. In contrast to the prior work on disentangled deep-learning [20-22], we propose a novel technique for interpreting the disentangled latent representations by mapping them to different classes of hierarchical spatial features, extracted from reconstructed images, through a process called visual latent traversals. (3) *Generalizability*: The strategy of gaining interpretability in MorphoGenie mimics the concept of combinatorial generalization in human intelligence that assemble different hierarchical spatial attributes from diverse image data to learn the unseen image data - thus facilitating the discovery of biologically meaningful inferences, especially the heterogeneities of cell types and lineages. Indeed, MorphoGenie is widely adaptable across various imaging modalities and experimental conditions, promoting cross-study comparisons and reusable morphological profiling results. The model generalizes to unseen single-cell datasets and different imaging modalities while providing explanations for its predictions. Overall, MorphoGenie could spearhead new strategies for conducting comprehensive morphological profiling and make biologically meaningful discoveries across a wide range of imaging modalities.

## RESULTS

### Overview of MorphoGenie

The learning part of MorphoGenie employs a hybrid neural network architecture, enabling the generation of a "disentangled representation" within its latent space — a model’s internal, high-dimensional conceptualization of data — while also facilitating the reconstruction of high-fidelity cellular images (**Fig. 1**). *Disentangled* representation is a concept that involves segregating and identifying independent variables (or factors of variation) that constitute the diversity observed within image data. In other words, a representation where a change in one latent dimension corresponds to a change in one factor of variation, while being relatively invariant to changes in other factors. In the context of cell morphology, these variables can be related to quantifiable attributes such as cell/nuclear size, texture of the cytoplasm, or specific cellular spatial patterns.

**Figure 1:**
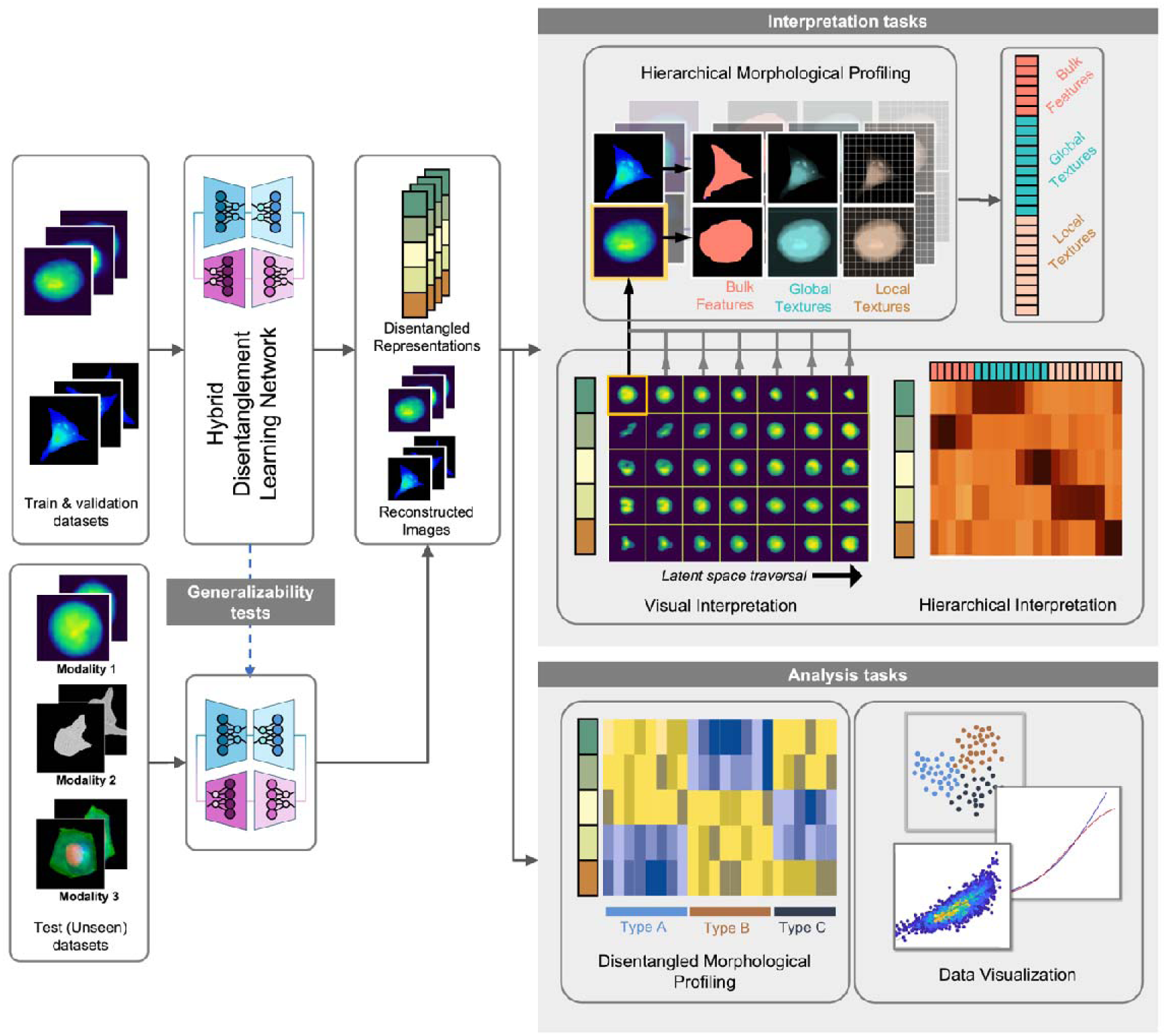
Overview of the MorphoGenie framework. It illustrates the sequential flow of tasks made possible through the integration of disentangled representation learning and high -fidelity image reconstructions. These tasks encompass morphological profiling and downstream analysis and the generation of interpretation heatmaps specific to the training dataset. Additionally, the figure highlights the utilization of a pretrained model, which facilitates cross-modality generalizability for morphological profiling and interpretability within the framework.

In MorphoGenie’s latent space, each disentangled dimension corresponds to one of these variations independently. Hence, it allows for visual *latent traversal*, a process that modifies only one chosen variable, with other factors remaining unchanged, thus enabling visual inspection of how individual features impact cell morphologies (**Fig. 1**). This capacity for selective alteration is crucial in deconvoluting the complexities of cellular morphology and understanding the distinct contributions of each morphological aspect.

Prior research has utilized VAEs for unsupervised learning in single-cell imaging, such as the use of VQ-VAE to predict cell state transitions [16, 23]. These approaches, however, often result in discrete and non-continuous latent spaces that lack the desired disentanglement, complicating the interpretation of morphological changes. Another work explored adversarial autoencoder (AAE) models for classification and identifying metastatic melanoma, but it, too, yielded entangled representations, impeding clear and direct downstream analysis [17].

These previous methodologies highlight the inherent trade-offs faced when seeking disentangled representations with VAEs. Techniques such as β-VAE and other factorized approaches have been proposed to enhance interpretability by creating a more structured latent space [20, 21]. However, the compact nature of these VAE-derived latent representations often leads to a loss in the quality of reconstructed images, presenting significant hurdles in mapping the disentangled latent factors to visually interpretable and biologically relevant features. In contrast, GANs excel in generating realistic reconstructions but typically result in a more entangled latent representation, which can obscure the direct interpretability required for precise morphological analysis. Also, GANs are known to suffer from training instability [24].

MorphoGenie is built upon bringing two generative models (VAE and GAN) into a hybrid architecture (**Fig. 2a**). The overall rationale is to jointly optimize the objectives of disentanglement learning and high-fidelity image generation by a dual-step training approach. Initially, a VAE variant, called FactorVAE (**Methods**), is employed to effectively learn the disentangled representations (in the latent space) from real image space, using a probabilistic encoder [20]. Subsequently, image reconstruction from the latent representation is accomplished through a decoder. During optimization, the objective is to minimize the disparity between the reconstructions and real images while at the same time learning the latent disentangled representations. FactorVAE is proven to provide a better trade-off between disentanglement and reconstruction performance than the state-of-the-art VAE models, notably the popular β-VAE [21]. In the second step, the disentangled representation learned from the first step is transferred to the GAN and trained by the generator to generate synthetic images, which are then assessed by a discriminator for differentiating between the generated (fake) and real images. By transferring the inference model, which provides a disentangled latent distribution, rather than a commonly used simple Gaussian prior, to GAN, this joint sequential learning approach could allow the overall hybrid model to learn latent representation by first learning the main disentangled factors by VAE, then learning additional (entangled) nuisance factors by GAN. Also, it allows a more accurate and detailed reconstruction than VAEs alone can achieve (See **Methods**).

**Figure 2.**
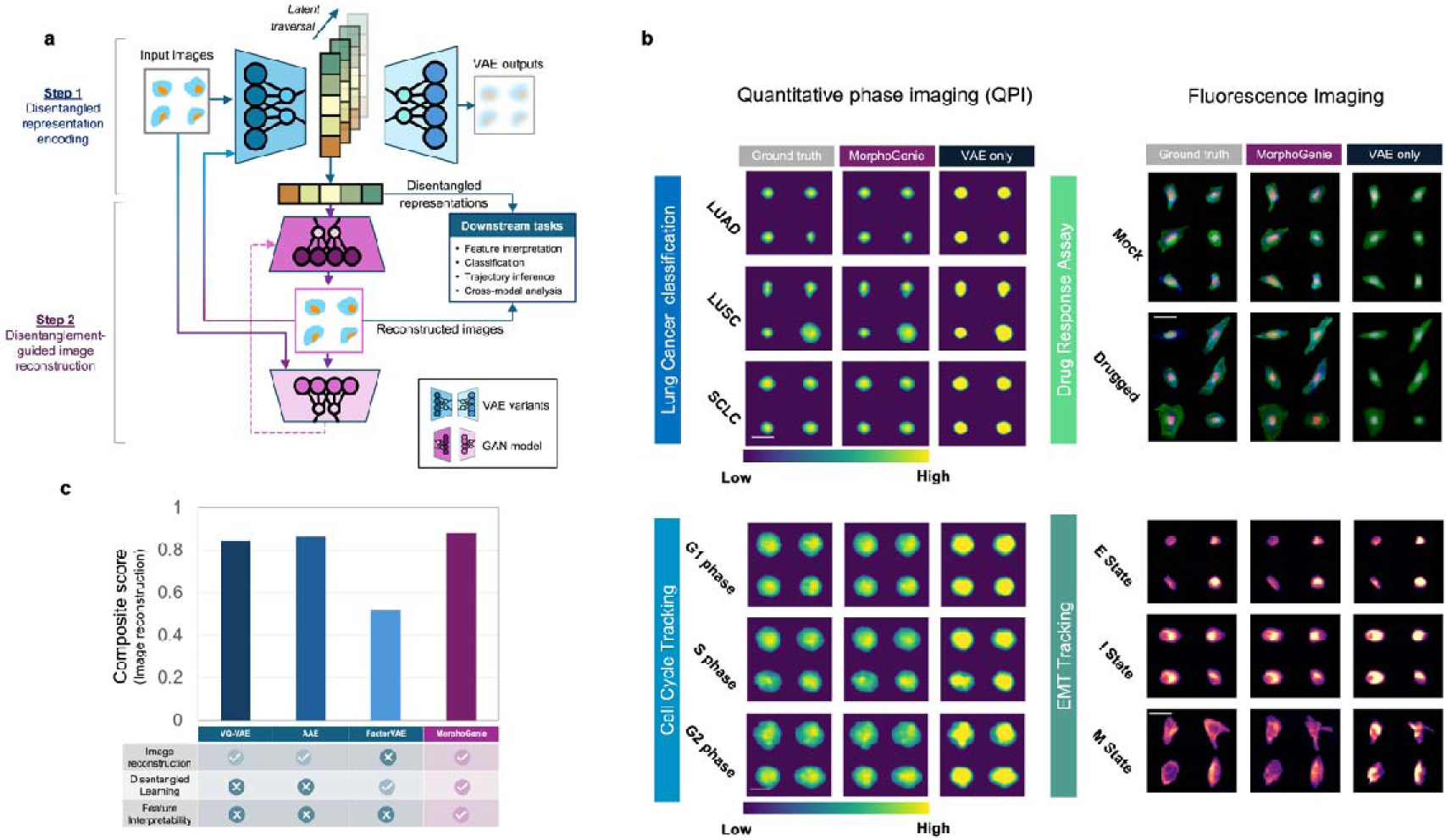
**a. Hybrid disentanglement learning network architecture in MorphoGenie.** It employs a dual-step learning strategy with an overall rationale that jointly optimizes the objectives of both the disentanglement learning and high-fidelity image generation. First, a VAE variant, called FactorVAE is employed to learn the disentangled representations in the latent space using a probabilistic encoder. Image reconstruction from the latent representation is accomplished through a decoder. In the second step, the disentangled representation is transferred to the GAN and trained by the generator to generate synthetic images, which are then assessed by a discriminator for differentiating between the generated and real images. This joint sequential learning approach could allow the overall hybrid model to learn how to minimize the disparity between the reconstructions and real images while at the same time learning the latent disentangled representations. **b. Image reconstruction performance in MorphoGenie in four distinctively different cell image datasets:** Quantitative phase images (QPI) of suspension cells (lung cancer cell type classification and cell-cycle progression assay, scale bar=20μm) and fluorescence images of adherent cells (Cell-Painting drug assay (scale bar=65μm) and epithelial-to-mesenchymal transition (EMT) assay, scale bar=30 μm). The lung cancer cell image datasets include three major histologically differentiated subtypes of lung cancer i.e., adenocarcinoma (LUAD: H1975), squamous cell carcinoma cell lines (LUSC: H2170), small cell lung cancer cells (SCLC: H526) are included. The cell-cycle datasets described the classified cell cycle stages (G1, S and G2 phase) of human breast cancer cells (MDA-MB231), scale bar= 20μm. The Cell-Painting drug assay dataset includes the human osteosarcoma U2OS cell line with (drugged) and without (mock) the treatment of glucocorticoid receptor agonist. The EMT dataset includes the A549 cell line, labeled with endogenous vimentin–red fluorescent protein (VIM-RFP), at different states of EMT, i.e., epithelial (E), intermediate (I) and mesenchymal (M states)**. c. Comparative analysis of image reconstruction performance**. The analysis is based on a composite score taking into account Structural Similarity Index (SSIM), Mean Squared Error (MSE) and Fréchet Inception Distance (FID) (**Supplementary Fig. 1**, and see **Methods**).

Prior work has highlighted that the disentangled factors could, to a certain extent, support combinatorial generalization - the ability to understand and generate novel combinations of familiar elements — a core attribute of human intelligence [25]. The idea is to capture the compositional information of the images through disentangled representation learning, and reuse (and recombine) a finite set of these representations to generate a novel understanding of the image in different scenarios (e.g., different imaging modalities as demonstrated in this work), thereby bridging the gap between human and machine intelligence [22, 26]. To this end, we attempt to systematically investigate the inter-relationship between the disentangled latent representation and the morphological descriptors of single cells extracted from a spatially hierarchical analysis (**Fig. 1**). Inspired by that the disentangled representations learned by some VAE variants have been shown effective for learning a hierarchy of abstract visual concepts [22], we define a hierarchy of single-cell morphological features segmented for this mapping analysis (using the classical mathematical/statistical metrics): from the fine-grained textures and their local multi-order moment statistics to the coarse-grained features such as cell body size, cell/organelle shape, cell mass density distribution and so on (**Fig. 1**). Based on this method, we establish a single-cell morphological profile in a hierarchy that allows us to gain semantic and biologically relevant interpretation of the disentangled representation - unraveling the factors governing single-cell generative attributes. This makes the model not only interpretable, as each latent dimension can be understood and visualized in isolation, but also transferable, as the learned representation can be applied to new, unseen data more effectively.

### Image reconstruction in MorphoGenie

We first assessed the performance of MorphoGenie in cell image reconstruction (**Fig. 2b**). High-quality image reconstruction in generative models is crucial for validating model accuracy, ensuring the encoded latent space captures and preserves the essential details of the input image data. Moreover, accurate image reconstructions allow users to make informed interpretability based on the model outputs. They also serve as a reliable reference for downstream applications, where the integrity of subsequent analyses depends heavily on the initial reconstruction’s fidelity. Importantly, we investigated how the hybrid architecture of VAE and GAN improves the reconstruction performance in MorphoGenie, compared to the case in which only VAE is included. We chose four distinctly different image datasets for the assessments, including quantitative phase images (QPI) of suspension cells and fluorescence images of adherent cells (**Fig. 2b**). In the case of multi-color fluorescence images captured by the Cell-Painting drug assays, in which morphologies of the key subcellular organelles can separately be analyzed, e.g., nucleus, mitochondria, nucleoli, actin, endoplasmic reticulum (ER)), we trained MorphoGenie with the multi-color image inputs, i.e. superimposing images of 5 fluorescence channels (i.e., dimensions 256x256x5) (See **Methods** for training details).

In general, the VAE-only model (based on FactorVAE) can only manage to preserve overall attributes such as shapes, sizes, and overall pixel intensities in both QPI and fluorescence images (**Fig. 2b**). However, the intricate local texture variations within cellular structures are generally lost. In contrast, MorphoGenie, which involves dual-stage training based on both VAE and GAN, effectively preserves the subcellular textural details in both QPI and fluorescence imaging scenarios (**Fig. 2b**).

We also compared MorphoGenie’s image reconstruction performance with the state-of-the-art models (VQ-VAE [23] and AAE [27]) adopted for cellular morphological analysis on the four different cell image datasets, i.e., covering diverse complex morphologies (both in suspension and adherent cell formats) and different imaging modalities (QPI and fluorescence imaging) (See the description of datasets in **Methods**) (**Fig. 2c**). To facilitate disentangled representation learning that can be easily interpretable (as discussed later), the latent space in MorphoGenie is kept to have only 10 dimensions, significantly smaller than the AAE (512 dimensions) [27] and VQ-VAE (1024 dimensions) [23] used in the prior work. Intriguingly, even with such a smaller dimensionality, MorphoGenie’s image reconstruction could still outperform these methods for the majority of datasets, based on a composite set of metrics, including Structural Similarity Index (SSIM), Mean Squared Error (MSE) and Fréchet Inception Distance (FID) values (**Fig. 2c**, **Supplementary Figure S1**).

To validate the importance as well as the robustness of the dual-stage training in ensuring high-fidelity image reconstruction, we further compared the reconstruction performance of the FactorVAE-only model and MorphoGenie across a wide range of a hyperparameter *γ* used in training for improving the representation disentanglement (**Methods**). Clearly, not only can MorphoGenie consistently demonstrate significant improvement in reconstruction performance compared to the FactorVAE-only model, but also exhibit relatively more stable image reconstruction performance (in terms of SSIM, MSE, and FID) across two orders-of-magnitude of *γ* (**Supplementary Fig. S2**).

### Disentangled representation learning and interpretation in MorphoGenie

Accurate image reconstruction by generative models is a complex task that does not inherently lead to disentangled and interpretable latent representations. MorphoGenie is designed to produce such disentangled representations (e.g., D1 and D2 in **Fig. 3a**), enabling the isolation and visual identification of key factors of variation in cell morphology. As mentioned earlier, the value in each disentangled dimension, which corresponds to one of these variations, is varied (e.g., vary D1), with other factors remaining unchanged (e.g., fix D2) - a process called *latent traversal*. Using MorphoGenie’s decoder to reconstruct the image, one can further perform visual inspection of how changes in individual dimensions (latent traversal) impact cell morphologies (**Fig. 3a**).

**Figure 3.**
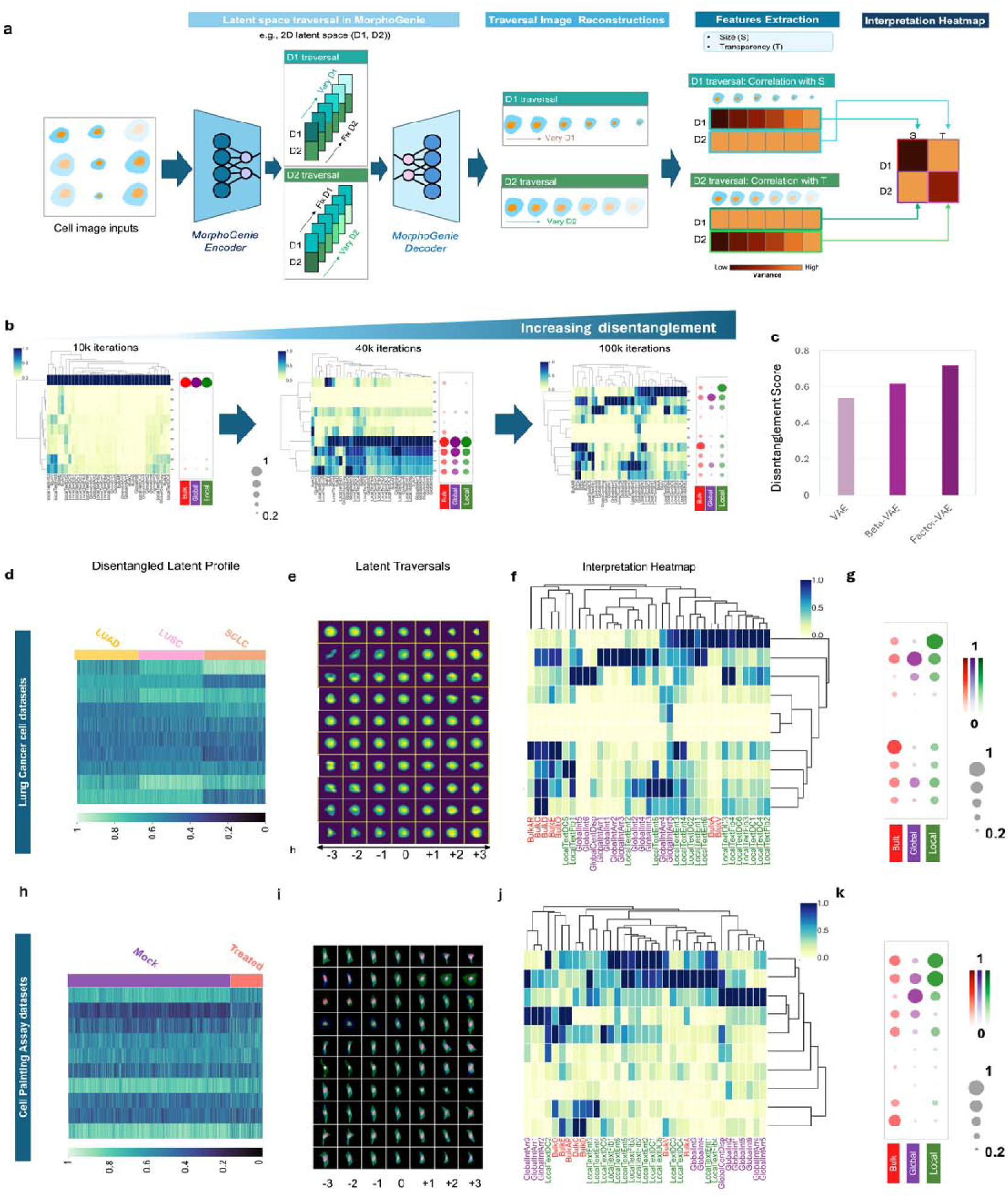
Disentangled representation learning in MorphoGenie. **a**. The workflow for generation of the ‘interpretation heatmap’. Consider a MorphoGenie model that generates a 2-dimensional latent space (D1, D2) only, we can visually assess how D1 (or D2) traversal (i.e., changing the D1 (or D2) disentangled representations while keeping D2 (or D1) fixed) impact the reconstructed images. Then, we can further investigate how the D1 and D2 traversals respectively correlate with different manually extracted features (e.g., cell size (S), and cell transparency (or opacity) (T)). Based on the statistical variance for each of the 2 features across these M (= 6 in this example) reconstructed images in the traversal, one could summarize a 1 x 2 vector that corresponds to the variance values of 2 features for a single latent dimension. This process is repeated for all other latent dimensions, generating 2 such 1x2 vectors, which are combined to create a 2 x 2 matrix (See M**ethods**). **b**. Using the interpretation heatmap together with the bubble plots, we show how the MorphoGenie model progressively learn to disentangle the bulk, global, and local morphological features across different latent dimensions throughout the training process (with an increasing number of training iterations using the lung cancer cell datasets). **c.** Disentanglement scores of different VAE models (vanilla VAE, and β-VAE and MorphoGenie based on FactorVAE). Figures **d-g** and **h-k** illustrate the interpretation pipeline, encompassing several key stages: **d, h** Disentangled latent (morphological) profiling to reveal the heterogeneities within the cell population, uncovering the existence of distinct types and subtypes. **e,i**. Latent space traversals. These traversals offer a qualitative visual insight into the variations of cellular features across different latent dimensions. **f,j** Interpretation heatmaps. **g,k.** Summarized bubble plot: This plot serves as a summary of disentanglement of the key cellular feature categories such as bulk, global and local. The interpretation pipeline is applied to the (**d-g)** lung cancer cell datasets (small cell lung carcinoma (SCLC), squamous cell carcinoma (LUSC) and adenocarcinoma (LUAD)) and (**h-k**) the Cell-Painting assay dataset (Human osteosarcoma U2OS cell line treated with glucocorticoid receptor agonist).

It should be noted that our work does not aim to benchmark various existing disentanglement metrics, among which no single disentanglement standard has proven consistently applicable across varied data types [20, 21, 28, 29]. Instead, we focus on evaluating the separability of latent features in relation to three key *visual hierarchical concepts/primitives*: *bulk, global textures, and local textures* **(Fig. 1)**. Bulk-level features primarily focus on cell size, shape, and deformation, while global texture features are based on the holistic textural characteristics of pixel intensities using multi-order moment statistics. Local texture features are extracted using spatial texture filters at various kernel sizes, quantifying local textural characteristics at both coarse and fine scales [10, 30, 31]. This approach is motivated by the demonstrated effectiveness of VAE models in learning hierarchical abstractions of visual concepts [22]

In order to assess how these hierarchical features can be discerned within the latent space, we introduce an *interpretation heatmap* that correlates the variability of hierarchical single-cell morphological features with that in the latent space dimensions (e.g., D1 is highly correlated with cell size changes shown in **Fig. 3a).** Based on the categorization of the three hierarchical primitives, we further devised a *disentanglement scoring system* (from the interpretation heatmap) that rewards higher separability of factors across different latent dimensions and penalizes scenarios with coexisting or entangled features within the same latent dimension (**Methods**). The disentanglement across the three hierarchical primitive factors can further be summarized by a bubble plot (**Fig. 3b**), in which we can gain further insights into the separability of the three visual primitive factors across the latent dimensions by highlighting the predominant factors of variation encoded in MorphoGenie. Thus, one can evaluate the extent of entanglement/disentanglement between the bulk, local, and global factors within each dimension of the latent space (**Methods**).

The overall approach of disentanglement learning assessment also provides a practical visual guide for the selection of the best-disentangled model, which can be determined through a grid search, tuning a hyperparameter that controls the degree of disentanglement in VAEs (e.g., β-VAE and Factor-VAE) (**Fig. 3b**, **Supplementary Fig. S3**). In this case, the evolution of the disentanglement learning process can clearly be visualized by both the interpretation heatmap and bubble plot. In the initial training steps, the latent dimensions predominantly comprise entangled features (**Fig. 3b**). However, as the training progresses, there is a noticeable transition during which the latent space becomes increasingly disentangled. Comparing different VAE models based on the *disentanglement scores*, Factor-VAE is chosen in MorphoGenie because of its consistently superior disentanglement performance across diverse image datasets (**Fig. 3c and Supplementary Fig. S3-4**).

We next demonstrate the MorphoGenie’s ability to learn the disentangled representations in two distinctly different image contrasts and cell formats - *label-free QPI* of three different lung cancer cell subtypes in *suspension* (**Fig. 3d-g**); and *fluorescence adherent* cell images of a Cell-Painting drug assay [12] (**Fig. 3h-k**). The disentangled latent profiles generated by MorphoGenie show distinct groups associated with specific cell types (i.e., small cell lung carcinoma (SCLC), squamous cell carcinoma (LUSC) and adenocarcinoma (LUAD)) or the presence of drug treatment (Human osteosarcoma U2OS cell line treated with glucocorticoid receptor agonist) (**Fig. 3d, 3h).**

Based on these profiles, we generated the latent traversal maps from which we can identify qualitatively the key factors of variations in the latent space (**Figure 3e, i**). For instance, in the lung cancer cell dataset, Dimensions 0, 1, and 3 show distinct changes in the textural distribution of phase. In contrast, Dimension 7 displays a significant shift in overall cell shape (**Fig. 3e**). In the Cell Painting dataset, Dimension 4 exhibits a noticeable change in overall cell size/shape. In contrast, Dimension 7 contributes to drastic textural modification within the cell (**Fig. 3i**).

To investigate further how MorphoGenie learns to disentangle cell morphological features, we employed the interpretation heatmap to illustrate the grouping/clustering of specific hierarchical visual primitive features corresponding to the disentangled latent dimensions. In both datasets, hierarchical features can effectively be segregated into different latent dimensions with minimal redundancy - indicating that the latent representations are disentangled according to the spatial hierarchy (**Fig. 3f, j**). Specifically, we observed that Dimensions 0, 3, and 7 in the lung cancer datasets, which are among the top-ranked latent features for classifying the three cell types (**Supplementary Fig. S6a**), are primarily linked to the local texture, global texture and bulk mass/optical density features, respectively (**Fig. 3f**).

Similarly, in the Cell-Painting dataset, predominant texture feature variation is observed, with Dimensions 1 and 5 emphasizing more on the local textural aspects and Dimension 7 highlighting global textural features. On the other hand, Dimensions 3 and 4 are more related to the bulk cell shape (**Fig. 3f and j**). Hence, these observations suggest that the most significant morphological features sensitive to drug perturbation are closely linked to the local and global textural changes, whereas the bulk aspects are comparatively less significant.

We note that only some latent features always display clear factors of variations in the 10-dimensional latent space. For instance, Dimensions 4 and 9 in the lung cancer dataset, the minor important features for cell-type classification (**Fig. 3e**), show no apparent morphological variability in the latent traversal maps. This observation is consistent with the latent feature ranking analysis (**Supplementary Fig. S5**) and thus their negligible contribution to the interpretation heatmap **(Fig. 3f)** and the bubble plot **(Fig. 3g)**. Based on these analyses, we can disentangle the relevant morphological features learned by MorphoGenie, and further provide an interpretable assessment of their contributions to classification tasks (which are further detailed in the next section), based on the three hierarchical visual primitives.

We note that all the components in our interpretation pipeline, from the latent traversal reconstructions, the disentangled latent profile, the interpretation heatmap, the features ranking, to the bubble plot, are arranged/aligned in the same order of latent dimensions (**Fig. 3d-g** and **3h-k**). This alignment allows feature interpretation to be easily approached in a bi-directional manner. Firstly, the user can select dimensions based on qualitative assessment and investigate deeper into the profiles and corresponding interpretation heatmaps. Alternatively, they can pick the targeted disentangled dimensions, assess the dimensions based on the feature rankings, and visually identify the morphologically varying factors learned by MorphoGenie.

### MorphoGenie enables interpretable downstream analysis: cell type/state classification

Previously, we demonstrated that a biophysical phenotyping method based on a deep neural network model, which was trained with the manually extracted hierarchical morphological features, could effectively identify three distinct histologically differentiated subtypes of lung cancers [11]. In contrast to this supervised learning method, we evaluate whether the unsupervised disentangled representation learning in MorphoGenie can also support such downstream biophysical image-based analyses [10].

Therefore, we performed dimensionality reduction using the uniform manifold approximation and projection (UMAP) algorithm based on the set of MorphoGenie’s disentangled representations (i.e., the profile shown in **Fig. 3d**) learned from all single-cell lung cancer QPI. Visualizing the MorphoGenie’s latent space in UMAP reveals the clear clustering of the three major lung cancer subtypes based on their label-free biophysical morphologies: LUSC, (H2170), LUAD (H1975), and SCLC (H526). (**Fig. 4a**). Using these disentangled representations to train a decision-tree-based classifier (**Methods**) also yields high label-free classification accuracies among three cell types (80%-95%) (**Fig. 4b**). We note that even the dimensionality of MorphoGenie is kept at 10 dimensions only, which displays high degree of disentanglement (compared to higher dimensional latent space (**Supplementary Fig. S6**), significantly smaller than the AAE (512-dimensional) [17] and VQ-VAE (1024-dimensional) [16], MorphoGenie is still able to cluster clearly in UMAP different lung cancer cell types (**Fig. 4a**). In terms of classification tasks, we also note that MorphoGenie consistently outperforms other VAE-only models across all the datasets studied in this work **(Supplementary Figure S7-8)**. Hence, it further substantiates the advantages of MorphoGenie over other state-of-the-art VAE models from the perspectives of image reconstructions and downstream analysis, with the extra benefit of having disentangled representation.

**Figure 4:**
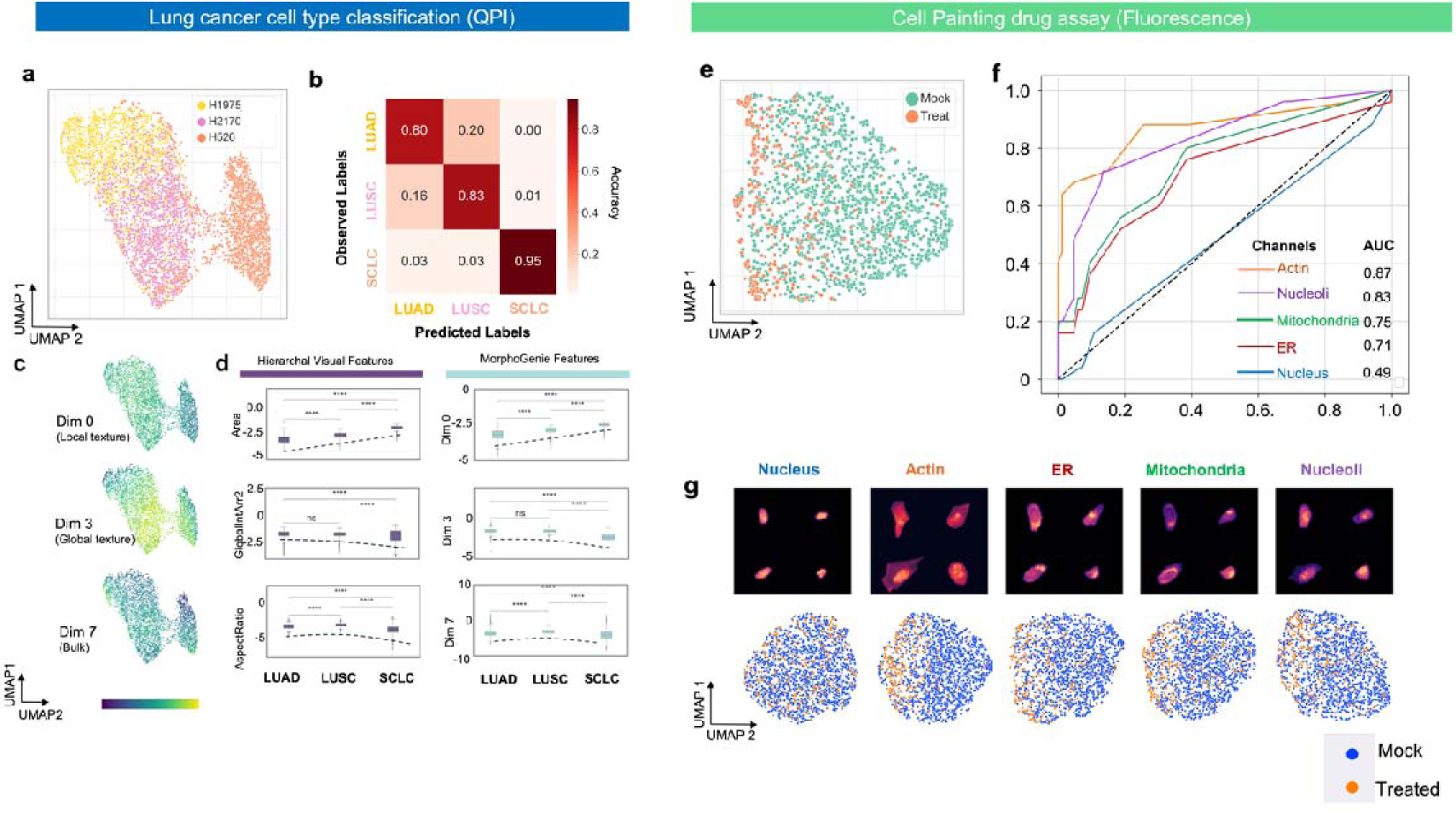
Downstream morphological analysis based on the disentangled representation learnt from MorphoGenie. **a-d. Analysis of biophysical morphology of lung cancer cell types LUSC, (H2170), LUAD (H1975), and SCLC (H526). a.** UMAP visualization of clusters of three lung cancer cell types. b. Label-free lung cancer cell type classification displayed in a confusion matrix. c. UMAP visualization (same as a) color-coded with the values of different disentangled MorphoGenie features (Dimensions 0, 3, and 7, representing local mass/optical density textural, global mass/optical density textural and bulk properties of cells). **d**. Quantitative comparisons of the manually extracted (hierarchical visual) features with the MorphoGenie features in three different aspects: bulk, global textures, and local textures. (**** represents p value < 1×10^-4^; ns: not significant, p-values reported from T-test with Bonferroni correction). **e-g. Organelle-specific analysis of fluorescence morphology of human osteosarcoma U2OS cell line treated with glucocorticoid receptor agonist. e.** UMAP visualization showing the shift of the treated cells from the mock population. **f.** Receiver operating characteristic (ROC) analysis showing the classification performance of different models trained with 5 fluorescent channels respectively (actin, nucleoli, mitochondria, endoplasmic reticulum (ER) and nucleus). **g.** (Top) Reconstructed images of different fluorescence (organelle) channels based on MorphoGenie features. (Bottom) UMAP visualizations for treated and mock cells based on the MorphoGenie features learnt from 5 different fluorescence (organelle) channels.

Further analysis of disentangled representations shows that the latent dimensions particularly influential in lung cancer cell-type classification, such as Dimensions 0, 3, and 7 (i.e., the dimensions that are more related to the three different hierarchical visual primitives morphological aspects (**Fig. 3f-g**)), display heterogeneities when color-coded in UMAP, confirming their discriminative power (**Fig. 4c**). We further observed that the patterns of the expressions (i.e., the normalized values) of disentangled representations among the three subtypes align very well with the manually extracted biophysical features falling under the same hierarchical attributes. For instance, Dimension 3 (primarily attributable to global texture of mass/optical density, as shown in **Fig. 3f-g**) shows the same pattern of variation as the global textural feature GlobalInt3 (Intensity Skewness in the intensity histogram) (See the box-plot comparisons in **Fig. 4d**). This agreement suggests that MorphoGenie’s representations are not simply statistically significant but also readily interpretable in label-free lung cancer subtype classification.

Apart from label-free QPI, we proceeded to test the downstream analysis with the *fluorescence* cellular images (based on the Cell Painting dataset shown in **Fig. 3g-j**). Clearly, the disentangled representations learned from MorphoGenie can show the morphological change/shift due to the bioactive compound treatment (**Fig. 4e**). To further identify the organelle-specific morphological variations, we subsequently trained MorphoGenie with separate fluorescence channels (**Methods**). We observed that MorphoGenie is indeed able to delineate the impacts on different organelles’ morphologies due to the drug treatment (**Fig. 4f-g**). Our decision-tree classifier trained with the disentangled latent representations showed that actin and nucleoli have the most distinct difference in morphology (AUC: 0.87 for actin; 0.83 for nucleoli) as AUC values (**Fig. 4f**). We also observed that the shifts in the drug-treated cell population from the control condition (mock) is more pronounced in the cases of actin and nucleoli, compared to other organelles (**Fig. 4g**).

### MorphoGenie enables interpretable downstream analysis: cellular progression tracking

We next investigated if MorphoGenie could extend beyond static cell type/state classification to enable dynamic tracking of cellular progression. Morphological profiling of continuous cellular progressions and dynamics could provide valuable insights into deciphering complex cellular development, e.g. cell growth, differentiation, and the mechanisms behind various physiological and pathological conditions. Here we put MorphoGenie to the test, evaluating its capacity to monitor morphological changes during cellular events such as epithelial-to-mesenchymal transition (EMT) and cell-cycle progression (**Fig. 5**).

**Figure 5:**
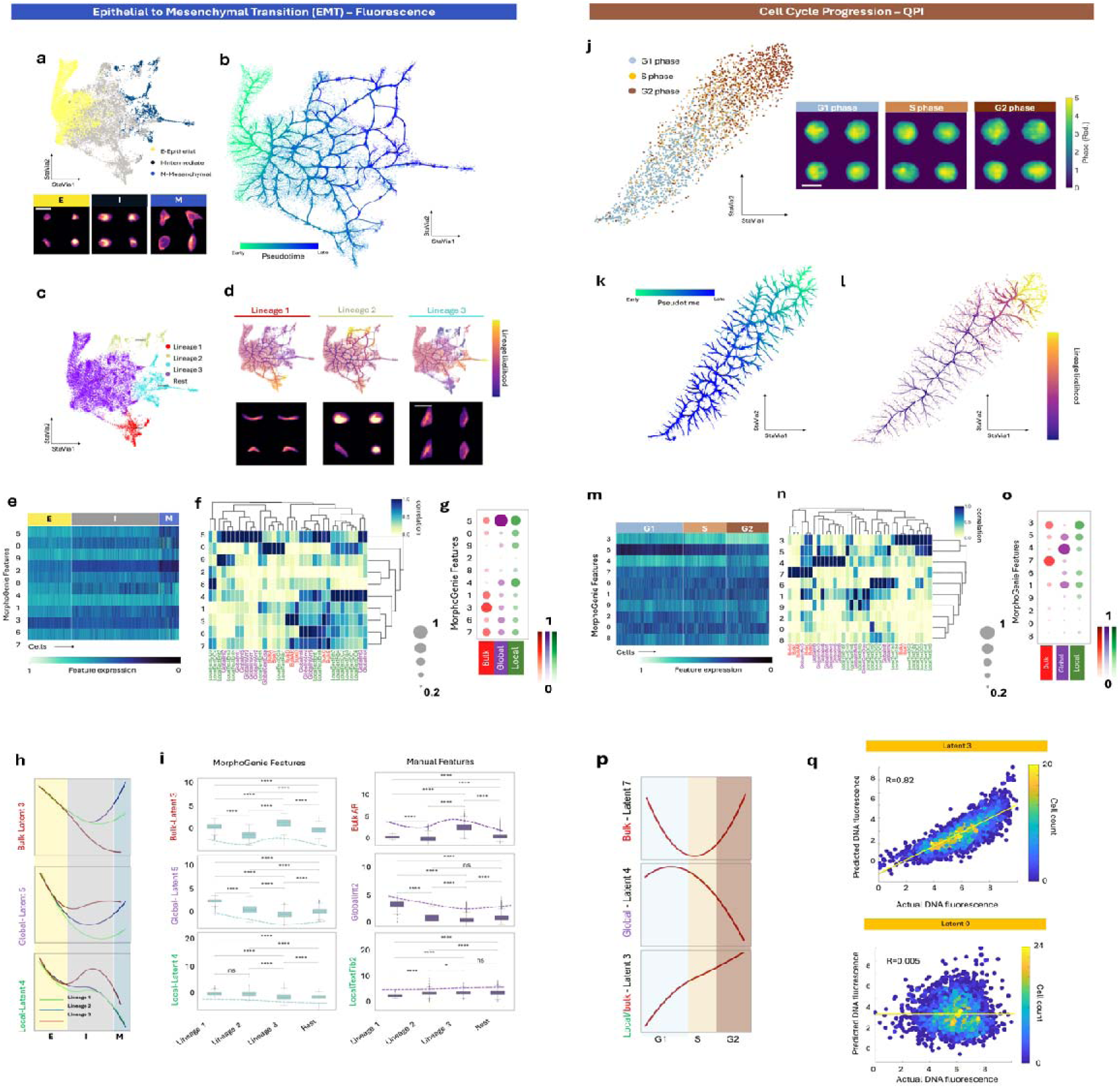
Downstream trajectory inference based on disentangled representations of MorphoGenie for (a-i) EMT tracking (fluorescence-labelled adherent cells) and (j-r) cell-cycle progression tracking (label-free suspension cells captured by QPI). **a. (Top)** Single-cell embedding visualization of EMT using StaVia based on the inputs from MorphoGenie’s features. **(Bottom)** Fluorescence images at three different stages of EMT: E, I, and M stages (scale bar = 30 um) annotated according the previous study [34]. **b**. An Atlas View of the pseudotime EMT trajectory of computed by StaVia, overlaid with a directional edge-bundle graph illustrating the overall pathway trend. **c.** Unsupervised lineage identification by StaVia. **d.** Color-coded lineage probability reveals and three terminal states and hence three pathways. Representative image of the morphologically distinct Lineages 1-3. (**e-g**) Feature interpretation pipeline for the EMT dataset: **e.** MorphoGenie’s disentangled morphological profiling. **f.** Interpretation heatmap. **g.** Bubble plot summary of the interpretation based on manually extracted hierarchical visual features. **h.** MorphoGenie feature trends in the pseudotime across E, I and M stages computed by StaVia (Dimensions 3, 5, and 4). **i.** Comparisons of the feature trends of the bulk, global and local factors in MorphoGenie with the corresponding manually extracted features (Dotted lines indicate the trend). p-value annotation legend: ns: not significant. *: 0.01 < p <= 0.05, **: 10^-3^ < p <= 0.01, ***: 10^-4^< p <= 10^-3^, ****: p <= 10^-4,^, p-values reported from T-test with Bonferroni correction. **j. (Left)** Single-cell embedding visualization of cell-cycle-progression (G1-S-G2 phase) using StaVia based on the inputs from MorphoGenie’s feature. The StaVia embedding is color-coded with the G1-, S- and S-phases, annotated independently by the fluorescently-labelled DNA images. (**Right**) Representative single cell QPI images of the cell states: G1, S, G2 (scale bar=20um). **k.** An Atlas View of the cell-cycle-progression, overlaid with a directional edge-bundle graph. **l.** Color-coded lineage probability showing the pathway G1-S-G2. **(m-o**) Feature interpretation pipeline for the cell-cycle dataset: **m**. MorphoGenie’s disentangled morphological profile. **n.** Interpretation heatmap. **o.** Bubble plot summary of the feature interpretation heatmap. **p.** MorphoGenie feature trends in the pseudotime across G1, S, and G2 stages, computed by StaVia (Dimensions 7, 4, and 3, representing the bulk, global texture and local texture features respectively). **q.** Correlative analysis between predicted DNA (by MorphoGenie) versus actual DNA content of cells in cell cycle progression (Higher correlation is shown in Dimension 3, compared to Dimension 0.

EMT is a critical process underlying various biological phenomena, including embryonic development, tissue regeneration, and cancer progression [32, 33]. EMT includes dynamic changes in cellular organization leading to functional changes in mobility. Here we utilized MorphoGenie to analyze time-lapse live-cell fluorescence imaging data from the A549 cell line, labeled with endogenous vimentin–red fluorescent protein (VIM-RFP). As vimentin, an intermediate filament, is a key mesenchymal marker, the expression and morphological changes of fluorescently-labeled vimentin could be indicative of the process of EMT, induced by transforming growth factor–β (TGF-β) in this study [34].

Furthermore, the disentangled representations derived from VIM-RFP-expressing cell images by MorphoGenie were scrutinized using a novel trajectory inference tool called StaVia [35]. It is an unsupervised graph-based algorithm that initially organizes single-cell data into a cluster graph [36, 37]. Subsequently, it applies a high-order probabilistic approach based on a random walk with memory to calculate pseudo-time trajectories, mapping out cellular pathways within the graph structure. StaVia also offers intuitive graph visualization “Atlas View” that simultaneously captures the nuanced details of cellular development at single-cell resolution and the overall connectivity of cell lineages in an edge-bundle graph format [37] (**Fig. 5b and k**). Through the integration of MorphoGenie and StaVia, we aim to provide a robust framework for providing holistic visual understanding of the continuous cellular processes with different complexities, as EMT and cell cycle progression studied in this work.

MorphoGenie’s disentangled representations (**Supplementary Fig. S9a**) successfully recapitulate chronology of the TGF-β-induced EMT into three different stages, i.e., epithelial (E), intermediate (I), and mesenchymal (M) (**Methods**) (**Fig. 5a**), annotated in the previous study [34]. Furthermore, MorphoGenie also reveals the heterogeneity in the single-cell EMT trajectories (**Fig. 5b**), in which three distinct EMT pathways leading to the separate terminal clusters (**Fig. 5c**). This is in contrast to the two pathways previously reported [34].

To validate this observation, we investigate the images of the mesenchymal populations in these three pathways, identifying discernible morphological differences (**Fig. 5d**). Particularly, we observe that the cells toward the end of pathway 2 and 3 (M) tend to show elongated spear shapes, characteristic changes of EMT [34]. This is compared to the cells at the terminal cluster of pathway 1 (I) are still in more or less round shapes.

Our trajectory inference analysis showcases differentially expressed disentangled representations (**Fig. 5e-g**) (notably Dimensions 3, 5, and 4) in the three pathways (**Fig. 5h**). We note that the representation trends in Pathway 3 are more deviated from the Pathway 1 and 2. Our interpretation heatmap highlights that Dimensions 3, 5, and 4 are more closely related to bulk, global texture, and local texture features of the cell morphology, respectively (**Fig. 5f, g**). On the other hand, our feature ranking suggests the dimensions that rank higher are pertaining to the shape-size morphologies and global intensity (**Supplementary Fig. S5c**). The above analyses thus provide a multifaceted description that suggests that the discriminative factors for EMT states are primarily associated with cell shape and size morphologies, and vimentin global textures, associated with subtle changes in local textures - consistent with the previous findings[34].

We further verified that the variations of the three disentangled representations (Dimension 3, 5 and 4) across three lineages correlate strongly with the corresponding hierarchical visual attributes (**Fig. 5i**). The distinct change patterns in Dimension 3 and 5 across lineages are highly consistent with the changes of aspect ratio (bulk AR) and the global intensity values (GlobalInt3-Skewness of the intensity levels of the image pixels). The small variation in Dimension 4 across lineages is concordant with similar insensitive changes in local texture features (LocalTextFib2-Radial distribution of fiber texture in the image).

We further explored MorphoGenie’s potential to predict cell cycle progression based on biophysical morphologies. For this purpose, we utilized our recently developed high-throughput imaging flow cytometer, FACED [38, 39]. FACED operates at speeds surpassing traditional imaging flow cytometry (IFC) by at least 100 times, offering a rapid and efficient alternative for cell cycle analysis, which typically relies solely on fluorescence intensity measurements from DNA dyes. Recent IFC advancements have demonstrated that label-free imaging can predict DNA content, and hence cell cycle phases, in live cells. MorphoGenie aims to expand on this capability by extracting biologically relevant information from large-scale single-cell FACED-QPI data and analyzing subcellular biophysical texture changes throughout the cell cycle (**Methods**).

In this study, our FACED platform captured a set of multimodal image contrasts, including fluorescence and QPI. These contrasts provide high-resolution insights into the biophysical properties of cells, such as mass and optical density, which are often challenging to assess through conventional methods. Utilizing QPI images from FACED, MorphoGenie generated disentangled representation profiles of live human breast cancer cells (MDA-MB231) as they progressed through the cell cycle (**Fig. 5j**, see latent traversal in **Supplementary Fig. S9b**).

Furthermore, using StaVia, our pseudotime analysis based on MorphoGenie’s disentangled representations reveals a well-defined progression (**Fig. 5j**), both in terms of pseudotime reconstruction (**Fig. 5k**) and lineage probability (**Fig. 5l**) - consistent with the chronology of G1, S and G2 phases, independently defined by the fluorescently-labeled DNA images in the same FACED imaging system (**Fig. 5j**, see **Methods**).

Based on our feature interpretation analysis (**Fig. 5m-o**), we can interpret that MorphoGenie’s disentangled representations captures general characteristics of changes in both cell size and local textures (Dimension 3), cell shape (Dimension 7), global textures (Dimension 4), composite changes in local/global textures (Dimensions 6 and 9) (**Fig. 5n-o**). In the pseudotime analysis, we indeed observed that Dimension 3, which captures simultaneous variability of cell size and local texture features, displays a significant progression through the G1-S-G2 phases (**Fig. 5p**). This finding aligns with established knowledge of cell growth in bulk size and mass during the cell cycle [38]. Moreover, Dimensions 4, 6 and 9, which are indicative of global/local phase intensity and relate to the dry mass density textures of cellular components like chromosomes and cytoskeletons [37], exhibit a slow-down during the G1/S transition (**Fig. 5p**) **(Supplementary Figure S10)**. This trend is in agreement with the slower rate of protein accumulation observed during the S phase [40].

This finding is further validated by the correlational analysis between the actual DNA content from the fluorescently labeled DNA images and the predicted DNA (**Fig. 5q**). In this analysis, Latent Dimension 3 shows a high correlation, with a pearson correlation coefficient of *R*=0.82, compared to other dimensions, which exhibit lower correlation coefficients. The dimension-wise correlation analysis is illustrated in **Supplementary Figure S11.** This suggests that dimension 3 is related to dry mass density growth relevant to cell-cycle progression. The strength of MorphoGenie, as demonstrated by these findings, lies in its ability to interpret the intricate biophysical morphology of cells captured by FACED and to provide a predictive analysis of cell cycle progression.

### Generalizability of MorphoGenie Across Imaging Modalities

The wealth of morphological data generated by current microscopy technologies poses a challenge for interoperable analysis, a key attribute for cross-modality correlative analysis, such as QPI-fluorescence imaging. For generative morphological profiling, achieving this interoperability necessitates a model with a high degree of generalizability. In the case of MorphoGenie, this means that the disentangled representations should encapsulate fundamental cellular image features and be transferable across diverse cellular image contrasts.

To assess MorphoGenie’s generalizability, we trained the model on a dataset from one imaging modality and tested its performance on unseen datasets with different image contrasts. We aimed to determine whether MorphoGenie could apply its trained latent representations to perform accurate downstream analyses and predictions on these new test datasets, without any retraining.

The robustness of a pre-trained MorphoGenie model is illustrated in Figure 6, where we demonstrate its ability to carry forward the learned insights as disentangled representations and make precise predictions in the unseen novel contexts. We evaluated four distinct models, each initially trained on a unique dataset. When tasked with analyzing three additional datasets, each with significant variations in single-cell analytical problems (from discrete cell state/type classification to continuous trajectory inference), cellular morphology (adherent versus suspension cells), and image contrasts (fluorescence, and QPI), the models produced visualizations with consistent global and local structures, whether viewed in UMAP or StaVia’s Atlas View (**Fig. 6a**)

**Figure 6:**
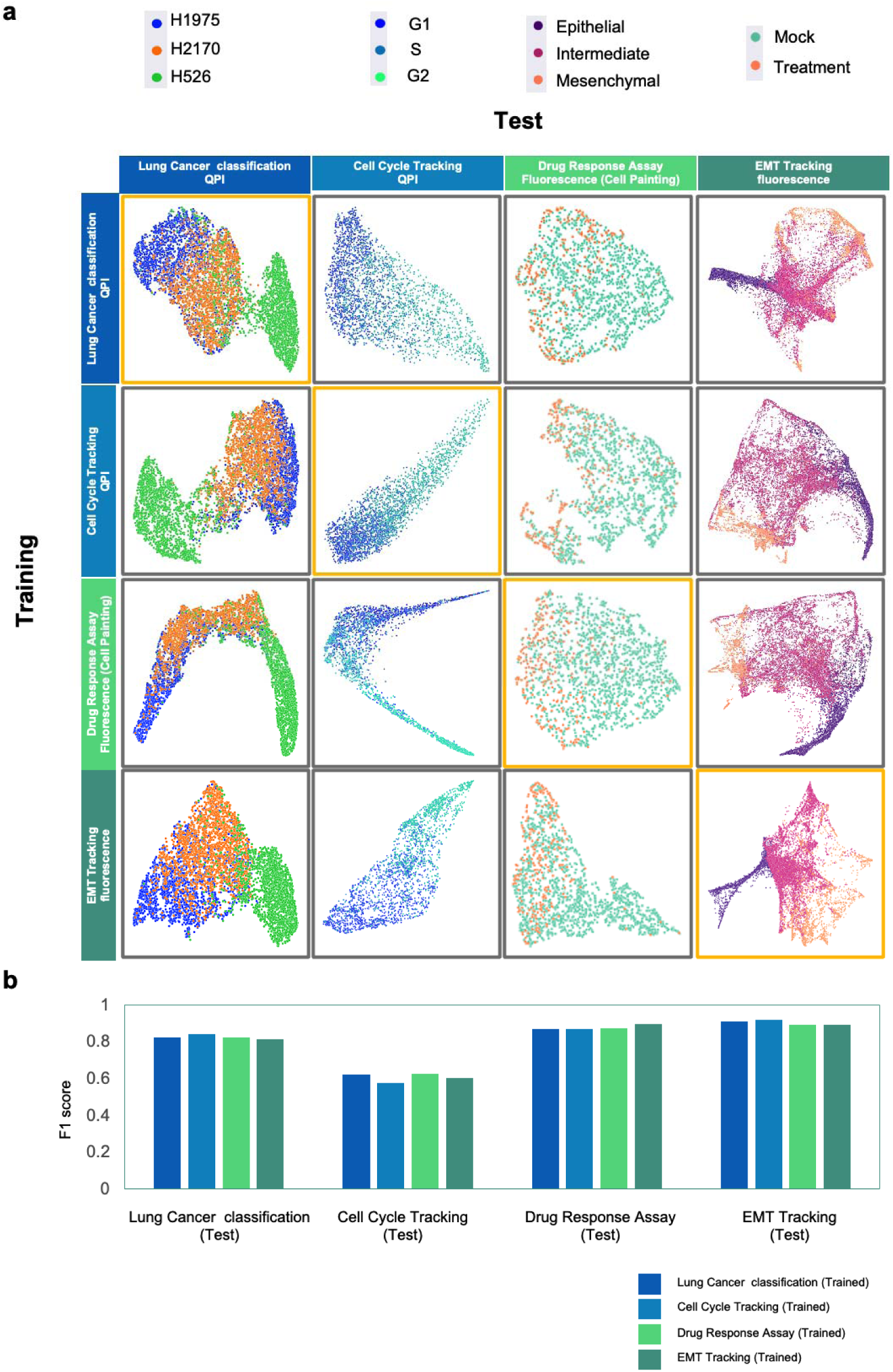
Generalizability performance of MorphoGenie. **a.** Qualitative 2D UMAP visualization. MorphoGenie is trained with one dataset (within orange box in each row) and tested on other datasets to predict the biological cell states and progressions. b. Quantitative assessments based on F1 scores for the generalizability tests across four datasets.

For example, a model pre-trained on the Cell-Painting assay dataset, which involved classifying morphological changes in adherent cells, was capable of classifying suspension lung cancer cells using label-free QPI images. Furthermore, it provided insightful trajectory inferences for cell cycle progression and EMT. The performance of such generalization, as evident from the F1 scores, is preserved across different model training scenarios (**Fig. 6b**).

The demonstrated generalizability of MorphoGenie reaffirms its value as an interoperable analysis tool, adept at handling the intricacies of multimodal imaging datasets. It is poised to play a pivotal role in integrating morphological profiling solutions, thereby enhancing our comprehension of cellular heterogeneity and dynamics across different imaging modalities.

## Discussion

Supercharged by the advances in computer vision, learning morphological features of cells through deep learning has gained considerable interest in the last decade. We introduced MorphoGenie, a deep-learning framework for profiling cell morphologies that effectively handles data from various microscopy modalities, including standard fluorescence, QPI, and imaging flow cytometry. Through comprehensive evaluations (from model performance to generalization), we demonstrated that MorphoGenie distinguishes itself from the current state-of-the-art with the following key attributes.

MorphoGenie is built upon a lightweight generative model that learns compact, interpretable, and transferable disentangled representations for single-cell morphological analysis. The model’s main strength is in extracting the compositional essence of cell morphology, distilling this into a limited set of key representations that can be flexibly interpreted to uncover novel insights across different imaging scenarios. This approach not only minimizes complexity but also enhances the alliance between human intuition and machine intelligence. Comparatively modest in dimensionality (**Supplementary Fig. S6**), MorphoGenie operates within a 10-dimensional latent space—significantly more concise than the expansive feature sets generated by manual extraction methods (e.g., Cell-Profiler: ∼1,700 dimensions [9]) or other generative models (e.g., AAE: 512 dimensions; and VQ-VAE: 1024 dimensions). Our evaluations illustrate its compatibility in a broad spectrum of single-cell image analyses, ranging from discrete cell-type classification to intricate trajectory inference. Compared to other state-of-the-art autoencoder models, MorphoGenie offers advantages of accurate image reconstruction (**Fig. 2, and Supplementary Fig. S2**), clear visual data analysis across diverse data types (**Supplementary Fig. S7**), disentanglement representation learning (**Supplementary Fig. S3, 4**, and **6**), and generalizable cross-modality learning (**Fig. 6**). MorphoGenie’s strength lies in its focused distillation of critical morphological information, avoiding the clutter of excessive features that can hinder meaningful interpretation.

MorphoGenie’s interpretability is further refined through a systematic investigation into the interplay between its disentangled latent space and the morphological descriptors of single cells, derived from a spatially hierarchical analysis. This analysis stratifies morphological features into a structured hierarchy from the nuanced textures (and their statistical analyses) to the more discernible attributes like cell size, shape, and mass/optical density textural distribution. Employing this hierarchical framework, MorphoGenie constructs a morphological profile that facilitates semantic and biological interpretations of the disentangled representations. It should be noted that it is not the purpose of MorphoGenie to seek for a universal interpretation that can offer a complete disentanglement among spatial hierarchical groups of cellular features - which is unlikely to occur in complex biological processes. Instead, MorphoGenie provides a practical reductionist framework to distill the key morphological features and their changes that can be explainable by the hierarchical features defined in this work. In fact, we argue that the interpretation can also be applied to any other types of features defined in different contexts (e.g., fractality [41], Fourier decomposition of cell shape[42] or other geometric and statistical descriptors [39] - as long as they could facilitate feature interpretation through MorphoGenie’s disentangled representation learning process.

Moreover, the generalizability of MorphoGenie is evidenced by its capacity to learn from one dataset and accurately predict morphological features in completely unseen datasets, regardless of the imaging modality, cell morphological formats and problems of interest. This adaptability demonstrated the model’s robustness and its potential as an interoperable analytical tool. It enables consistent cross-modality analysis, facilitating comparisons and integrations across studies, which is invaluable for the progression of biological research and understanding of cellular heterogeneity. MorphoGenie demonstrates flexibility in model selection, allowing for the integration of new generative algorithms that achieve disentanglement [43]. Incorporating these disentangled models with improved image generation quality enhances both interpretability and generalizability. This adaptability ensures that MorphoGenie remains at the forefront of technological advancements, continually improving its ability to extract meaningful insights from complex biological data.

Looking forward, there are a number of avenues for further development of MorphoGenie. (1) MorphoGenie’s capabilities can extend beyond individual cells. By learning factors that are not confined to hierarchical single-cell features, MorphoGenie can capture more fundamental attributes related to cell-cell interactions, protein localization, and other critical cellular processes. This expansion of scope enables the framework to provide deeper insights into the dynamics of multicellular systems, paving the way for broader applications in complex biological environments where understanding collective cellular behavior is essential. (2) 3D cellular imaging: Extending MorphoGenie to interpret three-dimensional cellular images will unlock a deeper comprehension of spatial cellular dynamics, benefiting from the volumetric data (e.g., confocal, multi-photon, and light-sheet imaging techniques). (3) Batch effect correction: Tackling batch effects could improve the model’s precision, minimizing technical noise and enhancing the biological signal in morphological data [44]. Leveraging the recent advancements in self-supervised/weakly supervised learning for this purpose will potentially improve the reproducibility and accuracy of phenotype classifications across different datasets. (4) Image reconstruction and translation: High-fidelity image reconstruction capabilities of MorphoGenie could augment the applicability of label-free imaging, such as translating QPI to fluorescence images. This could build a bridge between molecular specificity and morphological phenotypes, enriching our label-free understanding with detailed molecular insights. (5) Broadened interpretability: The framework currently categorizes features into bulk, global, and local textures. Expanding this taxonomy will capture a wider range of cellular features. It could be readily achievable in MorphoGenie, thanks to its ability to adapt to new domains where it can be fine-tuned to less-represented scenarios, potentially uncovering novel morphological features and contributing to the discovery of new cellular phenotypes or pathological states.

## Methods

### Disentangled Representation Learning in MorphoGenie

MorphoGenie is a deep-learning pipeline that generates the cellular morphological profiles and images of cells through disentanglement representation learning in an unsupervised manner. It can subsequently offer interpretation of the morphological profile through hierarchical feature mapping. Specifically, it employs a hybrid neural-network architecture built upon two generative models (VAE and GAN) (**Fig. 1-2**) that jointly optimizes the objectives of disentanglement learning and high-fidelity image generation by a dual-step training approach. In principle, while different VAE variants could be adopted in MorphoGenie, our comparative analyses have shown that FactorVAE stands out in accurately learning the disentangled representations in the latent space and reconstructing the cell images (**Fig. 3, Supplementary Fig. S2**). For the sake of clarity, we first describe the key features of different state-of-the-art VAE models tested in this work including the vanilla VAE, β-VAE and FactorVAE:

#### Variational autoencoder (VAE)

Consider a dataset consisting of *N* discrete or continuous variables x.

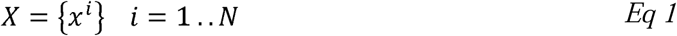

The encoder of a VAE initially maps the input data *X* to a probability distribution *q*_*e*_, which is modeled as a multivariate Gaussian distribution 𝒩 representing the latent space. The encoder learns to approximate the variables *z* of the *K*-dimensional latent space which is represented as a posterior approximation according to the Bayesian rule [15].

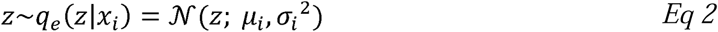

where *μ*_*i*_, *σ*_*i*_ (mean, variance) are the outputs of encoder. On the other hand, the decoder of the VAE samples the variable z from *z*∼*q*_*e*_(*x*_*i*_) to generate the observed data point x, which is given by

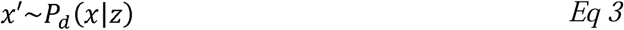

Assuming the data *X*′ is generated by continuous hidden representation z, by formulating a generative model Eq (4)

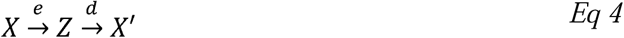

The above two approximations are optimized jointly by a single objective function:

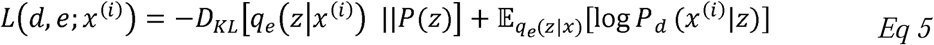

where *P*(*z*) is the prior which is a *K*-dimensional normal distribution 𝒩(0,1). The Kullback– Leibler divergence (KL divergence, *D*_*KL*_ [*q*_e_ (*z*|*x* ^(*i*)^) ||*P*(*z*)] is introduced to assess the divergence of the two distributions *q*_*e*_(*z*|*x* ^(*i*)^) and *P*(*z*). The above term of the LHS is optimized and differentiated to estimate the variational parameters ‘*e*’, and the generative parameters ‘d’. However, it is practically infeasible to estimate the parameter ‘*e*’ as it is not differentiable which is overcome by reparameterization trick. [15]

#### *β*-VAE

It extends the standard VAE by an additional hyperparameter β. β-VAE is deigned to achieve a disentangled latent representation by controlling beta. When β = 1 it represents a standard VAE and varying β>1 improves disentanglement at the cost of data reconstruction. However higher values of β allow interpretation of the latent space by varying dimensions, leading to an objective function [21]:

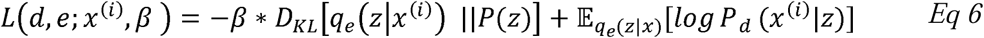

#### FactorVAE

FactorVAE (see **Supplementary Fig. S12)** addresses the tradeoff in β-VAE, in which penalizing the KL term with weight *β* encourages disentanglement at the same time. It leads to poor reconstruction as it reduces the amount of information of x in z. To address this, in FactorVAE, the KL term in the objective of VAE is split as

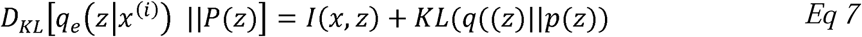

where, *I*(*x*, *z*) is the mutual information between the observation x and the latent variable z. However, penalizing the KL term in the above equation leads to pushing the posterior towards the factorized prior and, hence achieving a better disentangled independent latent factor. Therefore, the objective function of FactorVAE can be expressed as:

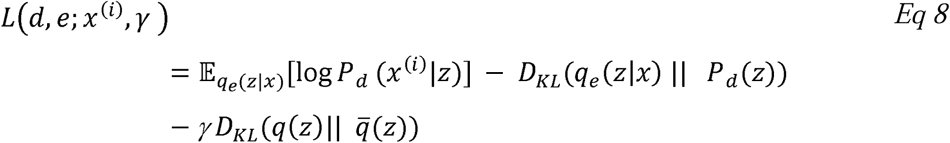

where,

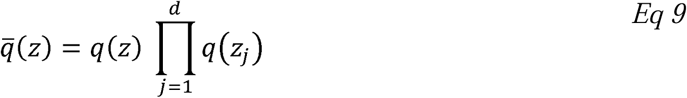

where *z*_*j*_ corresponds to one underlying factor of variation in the latent space. *γ* in Eq 8 directly encourages independence in the latent distribution and 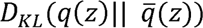 is known as a total correlation term, being intractable and is optimized by employing an alternate method called density-ratio trick, which involves training a classifier/discriminator **(Supplementary Figure S13)** to approximate the density ratio present in the KL term [20].

### Generative model in MorphoGenie

Disentanglement learning in VAEs often compromises the quality of generated images due to the strict constraints imposed to ensure that the latent variables are fully factorized (independent of each other). Our goal is to enhance VAE-based models by improving the quality of generated images while maintaining their ability to learn disentangled representations. To achieve this, we adopt a hybrid VAE-GAN architecture that separates the tasks of learning disentangled representations and generating realistic images into two distinct but sequential steps. First, we use FactorVAE to learn a disentangled representation of the data, ensuring that the latent variables are independent and capture distinct factors of variation. Next, we train a second network with a higher capacity for image generation by a GAN’s generator. This network takes the disentangled representation learned by the FactorVAE and decodes it into a high-quality, realistic image in the observation space. By decomposing these objectives, we leverage the strengths of both FactorVAEs and GANs: the FactorVAE focuses on disentangling the latent space, while the GAN enhances the quality of the generated images.

Our objective is to build a generative model (**Supplementary Figure S15**) that learns the generative parameter *ω* to produce high-fidelity output *x* with an interpretable latent code *z* (i.e. disentangled representation):

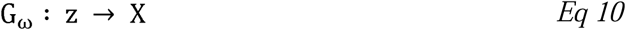

The idea is to decompose the objectives of the disentanglement learning and high-fidelity image generation as two different tasks [19]. Formally, let z = (*s*, *c*) denote the latent variable composed of the disentangled variable *c* and the nuisance variable *s* capturing independent and correlated factors of variation, respectively. Depending on which VAE model is used, the VAE model is first trained based on Eq. (5), (6) or (8) to learn disentangled latent representations of data, where each observation *x* can be projected to *c* by the learned encoder *q*_*e*_(*z*|*x*) after the training. Then in the second stage, encoder *q*_*e*_(*z*|*x*) is fixed to train a generator *G*_*ω*_(*z*) = *G*_*ω*_(*s*, *c*) for high-fidelity synthesis while distilling the learned disentanglement by optimizing the following objective:

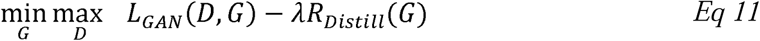

Aligning the reconstruction of GAN with the disentangled representation is achieved by maximizing the information between the disentangled latent representation and the generator output corresponding to the latent representation. *L*_*GAN*_ (*D*, *G*) is optimized in an adversarial manner for the discriminator to classify a real versus fake image; and is also optimized for the generator to improve image generation from the random noise:

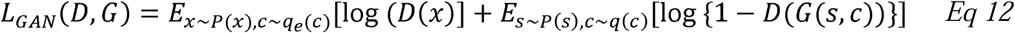

*R*_*Distill*_ term jointly maximizes the mutual information between the latent variable *c* and the generator output *I*(*c*; *G*(*s*, *c*)). Note that *c* is sampled from the aggregated posterior *q*_*e*_(*z*|*x*) instead of the prior *q*(*c*). And it is also optimized with respect to the generator only.

### MorphoGenie Training

The latent space dimension of the VAE models studied in this work is set as 10 (**Supplementary Fig. S6**). The encoder is trained with images of size 256x256x3. Encoder and decoder (**Supplementary Fig. S12**) and discriminator (**Supplementary Fig, S13**) of the FactorVAE is optimized using adam optimizer with decay parameters *β*_1_=0.9, *β*_2_=0.999 at a learning rate of 0.0001. On the other hand, the discriminator of the GAN in MorphoGenie is optimized with the decay parameters *β*_1_=0.5, *β*_2_=0.9 at a learning rate of 0.0001 and a batch size of 32. The generator of the GAN consisting of the resent blocks is trained with the latent vector (10-dimensional) and a random noise vector called as nuisance vector, which has 256 dimensions [19]. Generator and discriminator are trained at learning rate if 0.0001 using RMS prop optimizer, with a batch size of 32 (**Supplementary Fig, S14**).

### Model Selection

Model selection is performed by conducting a grid search to optimize the hyperparameters that control the strength of disentanglement in VAE, β-VAE, and FactorVAE. The best model is chosen based on its downstream performance and its ability to separate the three hierarchical primitive feature categories (bulk, global texture, and local texture) in the latent space. Subsequently, the top-performing disentangled models are further interpreted by generating the interpretation heat maps. It is important to note that the level of disentanglement varies among different the dataset and should be optimized by tuning the hyperparameter during model training.

### Cell Image Datasets

We choose datasets that are both open source and those imaged in-house to demonstrate the applicability of our approach to the datasets that are diverse in multiple aspects. Primarily, the choice of multiple imaging modalities and contrasts spanning fluorescence, bright-field and QPI. On the other hand, our choice of datasets also includes distinctly different biological conditions such as multiclass cell type classification (Lung cancer cell type (LC)) cellular response to drug treatment (Cell-Painting Assay (CPA)), and those showing continuous biological processes such as cell-cycle progression (CCy) and epithelial-to-mesenchymal transition (EMT). Further, inclusion of different imaging conditions, namely adherent cells (in EMT and CPA datasets) and cells in suspension (in Ccy and LC datasets).

#### Lung Cancer Cell-Types (LC)

This dataset consists of single-cell images (QPI) of three major histologically differentiated subtypes of lung cancer amongst seven cell lines, i.e., Three adenocarcinoma (H1975), squamous cell carcinoma cells (H2170), small cell lung cancer cell lines (H526). This data was collected by an in-house high-throughput microfluidic QPI flow cytometry system called multi-ATOM. Detailed experimental setup and protocols can be referred to ref. [11]. Using intensity-only measurements, multi-ATOM retrieves the complex-field information of light transmitting through the cell and yields two image contrasts at subcellular resolution: bright-field (BF: amplitude of the complex-field), and quantitative phase (i.e., QPI). BF images essentially display the distribution of light attenuation (or optical density) within the cell whereas QPI presents dry-mass density distribution within the cell [11].

#### Epithelial to Mesenchymal Transition (EMT)

This dataset consists of fluorescence images of adherent live cell image data (A549 cell line, labeled with endogenous vimentin–red fluorescent protein (VIM-RFP)) to study the morphological changes in response to TGF-β-induced EMT process [34]. In this dataset, the morphological dynamics of vimentin is quantified by extracting fluorescence texture features (Haralick features). The TGF-β treatment showed a shift in distribution for nearly all Haralick related features (i.e., texture features), with only two trajectories during EMT process were reported. A basic morphological operation has been performed in this work to annotate epithelial and mesenchymal cells by measuring the aspect ratio [34]. Elongated mesenchymal cell population is generally different from epithelial cells which are generally round and small. The remaining cells are categorized as the population in the intermediate state between the epithelial and mesenchymal states.

#### Cell Painting Assay (CPA)

This is a subset of BBBC022, which is a publicly available fluorescence Cell Painting image dataset, consisting of U2O2 cells treated with one of the 1600 bioactive compounds. In this dataset, images consisting of 5 channels tagged with 6 dyes characterizing 7 organelles (nucleus, golgi-complex, mitochondria, nucleoli, cytoplasm, actin, endoplasmic reticulum). The dataset is provided with annotation of the plate locations corresponding to the compound and the mechanism of action [12]. One of the treatments annotated as glucocorticoid receptor agonist named Clobetasol Propionate is used in this study for training and testing MorphoGenie. To test perturbation due to the chosen bioactive compound treatment, training is performed in two different ways. First, we overlaid multiple fluorescence channels of the images with the dimension of 256x256x *FL*, wherein *FL* can be more than 1 or can be extended up to the maximum number of fluorescence channels available in the dataset. Second, the models are trained separately by using the 5 different channels in the dataset.

#### Cell-Cycle Progression

This dataset was captured using another novel, in-house ultrafast QPI technique based on free-space angular-chirp-enhanced delay (FACED). It is an ultrafast laser-scanning technique that allows for high imaging speed at the scale orders of magnitude greater than the current technologies. More specifically, this FACED imaging system is integrated with a microfluidic flow cytometer platform enabling synchronized and co-registered single-cell QPI and fluorescence imaging at an imaging throughput of 77 000 cells/s with sub-cellular resolution [38, 39, 45]. This dataset was collected in an assay for cell-cycle progression tracking of human breast cancer cells (MDA-MB231) in microfluidic suspension. Annotations of the cell-cycle stages (G1, S, and G2) in this dataset were provided by quantitatively tracking the content of DNA labelled with Vybrant Dye orange stain (Invitrogen).

### Interpretation Heatmap

The latent space of MorphoGenie effectively encodes information of cell morphology within its disentangled dimensions. By traversing the latent space and reconstructing images from MorphoGenie, we can observe variations in the morphological features encoded within each latent dimension (**Fig. 1, 2a**). Furthermore, we can investigate how these latent features can be interpreted through a set of hierarchical cellular information. In detail, we manually defined and extracted a total of *M* = 35 morphological features from the latent traversal images, encompassing bulk, global textural, and local textural characteristics of cell morphology, i.e., *Feat* = {*feat*_1_,…. *feat*_*M*_}. The chosen latent space dimension is *K* = 10, which shows high degree of disentanglement (**Supplementary Fig. S6**). For each dimension, we computed *Q* (=10) steps of latent traversal and visualized the morphological variations across each traversal step. We calculated the statistical variance for each of the *M* (=35) features in each latent traversal of the *k*^th^ dimension, resulting in a 1 x *M* vector (**Fig. 3a**). This process is repeated for all *K* (=10) latent dimensions, generating *K* such 1 x *M* vectors. These vectors are then stacked to create a *K* x *M* matrix. We then performed hierarchical clustering in this matrix to identify the relationships between the *K* latent dimensions and *M* manually extracted morphological features in a heatmap representation. We refer to this clustered heatmap as the ‘*interpretation heatmap*’. This heatmap serves as a reference guide for our MorphoGenie feature analysis. For latent traversal of *K* = 10 latent dimensions and a total number of features extracted is *M* = 35, the *interpretation heatmap* (matrix) can be represented as

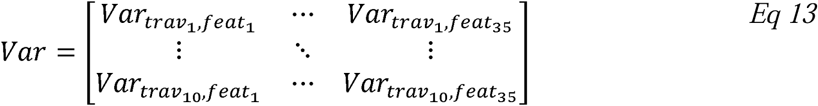

Var is a *K* x *M* (i.e., 10 x 35 in this example) matrix that is subjected to hierarchical clustering. Each element in the matrix (*Var*_*trav*_*k*_,*feat*_*m*__) is the variance of the feature (*feat*_*m*_) extracted from all Q steps across the *k*^th^ dimension traversal, which is computed as:

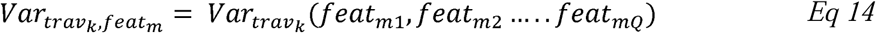

*feat*_*mi*_, where *i* = {1:*Q*}, is the feature value of *feat*_*m*_ extracted from the i^th^ traversal step. Based on these formulations, the interpretation heatmap provides valuable insights into the encoded features and their variations within the disentangled latent space, aiding our understanding of model predictions and generalization capabilities.

### Summarized Disentanglement Score

Various methods of defining and assessing disentanglement have been proposed in the previous studies [20, 21, 28, 29, 46]. Notably, β-VAE and FactorVAE metrics follow a supervised approach in which the annotations of the factors of variation in a dataset are predefined. However, in practical real-world datasets, whose annotations are not known, unsupervised disentanglement metrics are necessary. While there is no universal definition of disentanglement, we here investigate and interpret disentanglement of MorphoGenie’s latent representations broadly in the context of three hierarchical primitive categories (factors), i.e., bulk, global texture and local texture features.

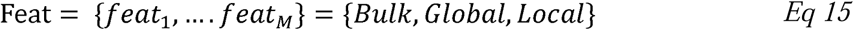

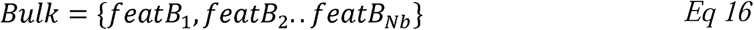

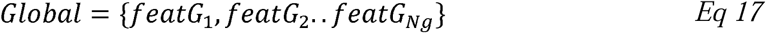

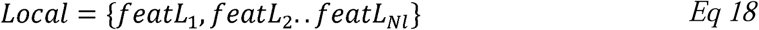

*N_b_*, *N_g_* and *N_l_* are the number of features in the hierarchical feature category of Bulk, Global and Local textures, respectively. Following Eq. (13), we can then define a matrix *S* that summarizes the significance of the hierarchical primitive factor/category in each latent dimension:

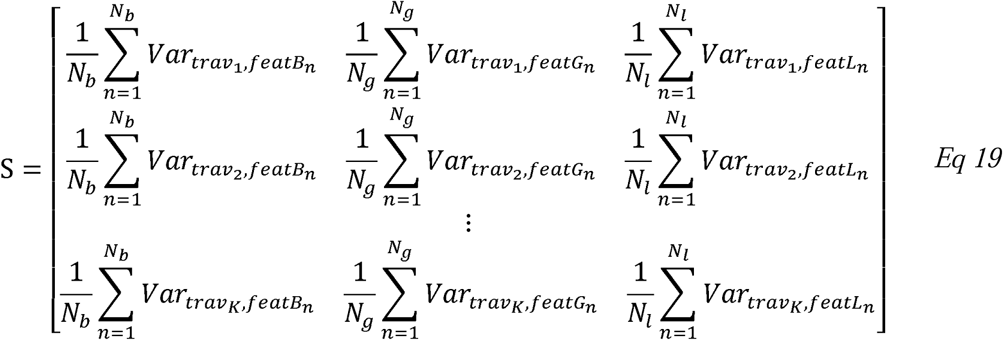

where K = 10 is the latent dimension in this work. In fact, the matrix S can be represented as the bubble plots shown in Fig. 3 and 5. This matrix computes, in each latent dimension, the mean of the variances of all the features groups in the three hierarchical primitive categories. The features exhibiting higher variance values spotlight the factors of variation tied to the latent dimension. This approach helps us understand the specific attributes that contribute significantly to the variations within the latent space. Hence, matrix S essentially displays the importance of each category in each latent dimension.

Based on this matrix S, we can compute a *disentanglement score* which reflects how the three hierarchical primitive categories can be separated to different latent dimensions. Specifically, the maximum value in each column (i.e., each hierarchical primitive category) corresponding to either bulk, global and local texture attributes in matrix *S*, i.e., *MaxVar*_*Bulk*_, Max*Var*_*Global*_, *MaxVar*_*Local*_ is regarded as the key bulk, global texture and local texture attributes respectively and the mean of the three variances is computed as the *disentanglement score*.

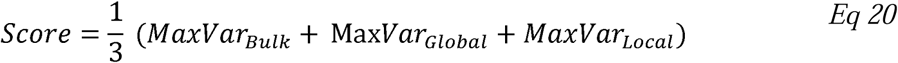

The model is said to be more entangled if two categories having maximum mean values in the same latent dimension, resulting in a lower disentanglement score.

### Other Performance Metrics

#### Mean Square Error (MSE)

MSE is a metric used to evaluate the accuracy of image reconstruction. It calculates the average of the squared differences between actual and predicted pixel values. In the context of MorphoGenie, *y* and 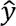 is the pixel values of the real image and the generated image. With *N* as total number of pixels in the image, MSE is computed as:

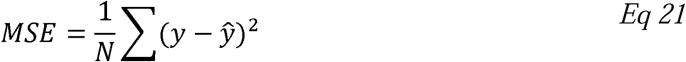

#### Fréchet inception distance (FID)

FID is a metric used to evaluate the quality of images generated by generative models, such as VAEs and GANs. It measures how similar the latent representations of generated images are to those of real images. Specifically, FID calculates the distance between these distributions, with lower values indicating closer resemblance and better performance of the model in generating realistic images.

#### Classification Accuracy

F1 score is used to measure the classification accuracy of the model based on the true positive (TP), false positive (FP) and false negative (FN) values from the confusion matrix of a tree-based decision classifier:

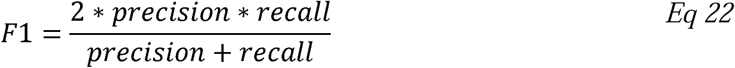

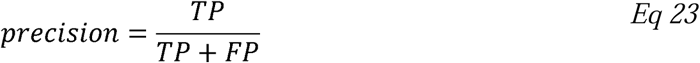

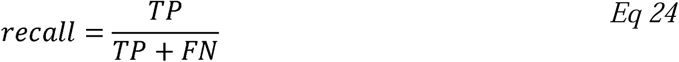

#### SSIM

SSIM is a metric used to evaluate the similarity between two images by analyzing their luminance, contrast, and structural characteristics. It ranges from 0 to 1, where 1 indicates perfect similarity. In this work, SSIM is applied to compare real and reconstructed images, averaging over 500 reconstructions, to assess the efficiency of the deep learning model in image reconstruction.

#### Feature Ranking

Importance of disentangled latent representations is measured by decision tree-based classifier. This basically works by computing how much impurity is reduced by each feature and hence determining the importance of every feature in classifying the samples according to given labels. Impurity here refers to presence of samples of one category under the label of another category.

### StaVia

StaVia is a computational framework for trajectory inference and visualization based on the diverse single-cell multi-omics as well as image data, including time-series data. StaVia uses higher-order random walks with teleportation and lazy-walk characteristics, which consider cells’ past states, to deduce cell fates, differentiation pathways, and gene (or morphological feature) trends, without imposing the assumptions about the underlying data structure. Built on the VIA framework [35], StaVia performs trajectory inference at both the single-cell and cluster-graph levels, handling complex non-tree topologies in large-scale datasets efficiently, even with millions of cells and modest computational resources. StaVia can also visualize inferred trajectories using an "Atlas View," a cartographic approach that provides intuitive graph visualization at large scale. This view captures detailed cellular development at single-cell resolution while also illustrating broader cell lineage connectivity, such as cell cycle progression and epithelial-mesenchymal transition (EMT).

## Code availability

MorphoGenie source code with tutorials to setup and test sample data is available on github. Link: https://github.com/rashmisrm/MorphoGenie

## Contributions

K.K.T. and R.S.M. conceived the project. R.S.M developed the algorithm and software to analyze the data. K.K.T. and R.S.M., S.V.S., and M.C.K.L. analyzed data. D.M.D.S. and G.G.K.Y. performed the imaging experiments and generated QPI datasets. R.S.M. and K.K.T. wrote the paper. All authors commented on and edited the text.

## Acknowledgements

The work is supported by Advanced Biomedical Instrumentation Center, the Research Grants Council and the Innovation and Technology Commission of the Hong Kong Special Administrative Region of China (grant nos. 17125121, 17208918, RFS2021 - 7S06, and ITS/318/22FP), Platform Technology Funding of the University of Hong Kong.

