## Supplementary Information for "Generalizable Morphological Profiling of Cells by Interpretable Unsupervised Learning"

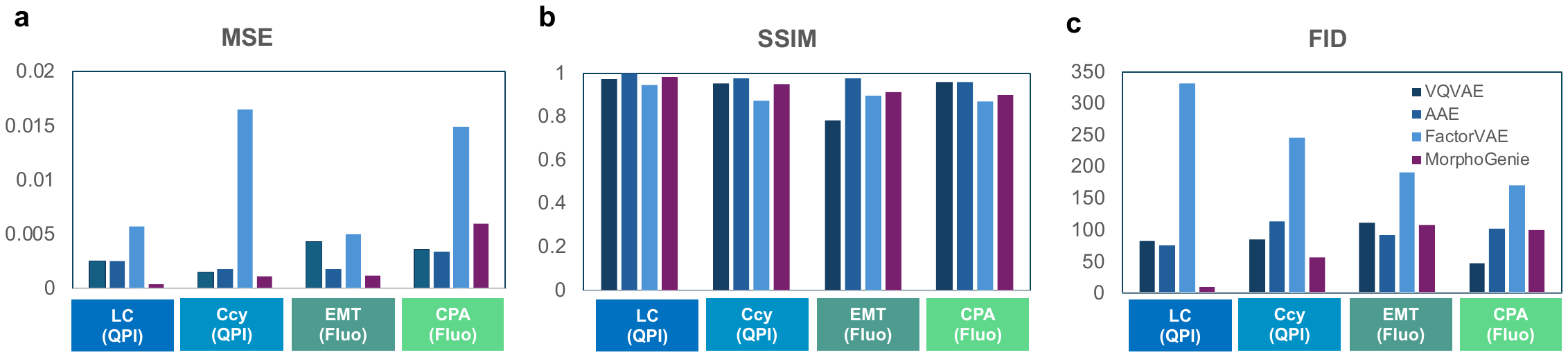

**Figure S1:** Comparisons of MorphoGenie against other state-of-the-art autoencoder methods (VQ-VAE, AAE, FactorVAE) for image reconstruction quality in terms of (a) Structural Similarity Index (SSIM), (b) Mean Squared Error (MSE) and (c) Fréchet Inception Distance (FID) values (See **Methods** of the definitions of SSIM, MSE and FID). Four different cell image datasets covering diverse complex morphologies (both in suspension and adherent cell formats) and different imaging modalities (QPI and fluorescence imaging) are employed: lung cancer type classification based on QPI (LC), cell-cycle progression assay based on QPI (Ccy), fluorescence epithelial-to-mesenchymal transition assay (EMT), and Cell-Painting drug assay (CPA).

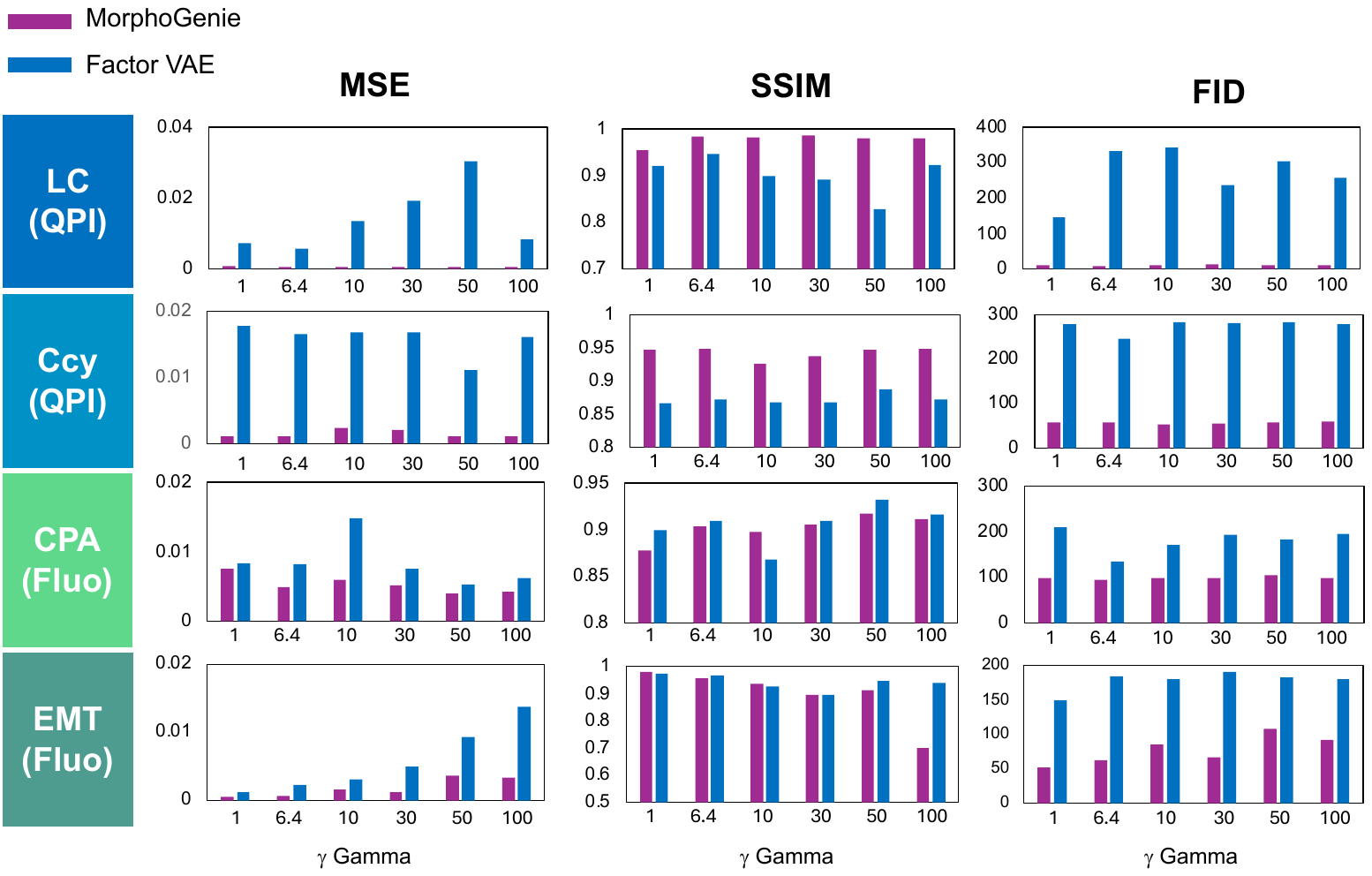

**Figure S2:** Comparative assessments of image reconstruction quality in MorphoGenie and FactorVAE-only model for a wide range of gamma $\gamma$, a hyperparameter used for improving representation disentanglement. Four different cell image datasets covering diverse complex morphologies (both in suspension and adherent cell formats) and different imaging modalities (QPI and fluorescence imaging (Fluo)) are employed: lung cancer type classification based on QPI (LC), cell-cycle progression assay based on QPI (Ccy), fluorescence epithelial-to-mesenchymal transition assay (EMT), and Cell-Painting drug assay (CPA).

**Figure S3:** Interpretation heatmaps generated using different methods to achieve disentanglement by adjusting hyperparameter strengths. A grid search method was employed to select the best disentangled model in MorphoGenie.

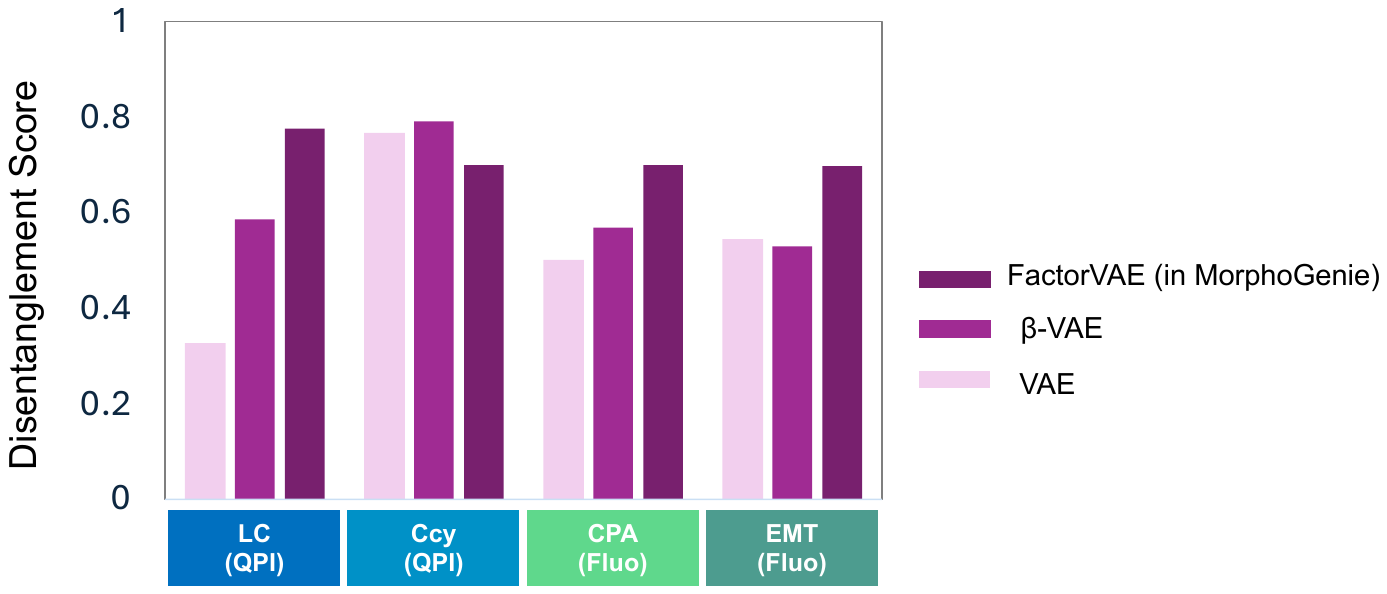

**Figure S4:** Disentanglement scores of vanilla VAE model, β-VAE model and MorphoGenie based on FactorVAE (on the datasets of LC, Ccy, EMT and CPA).

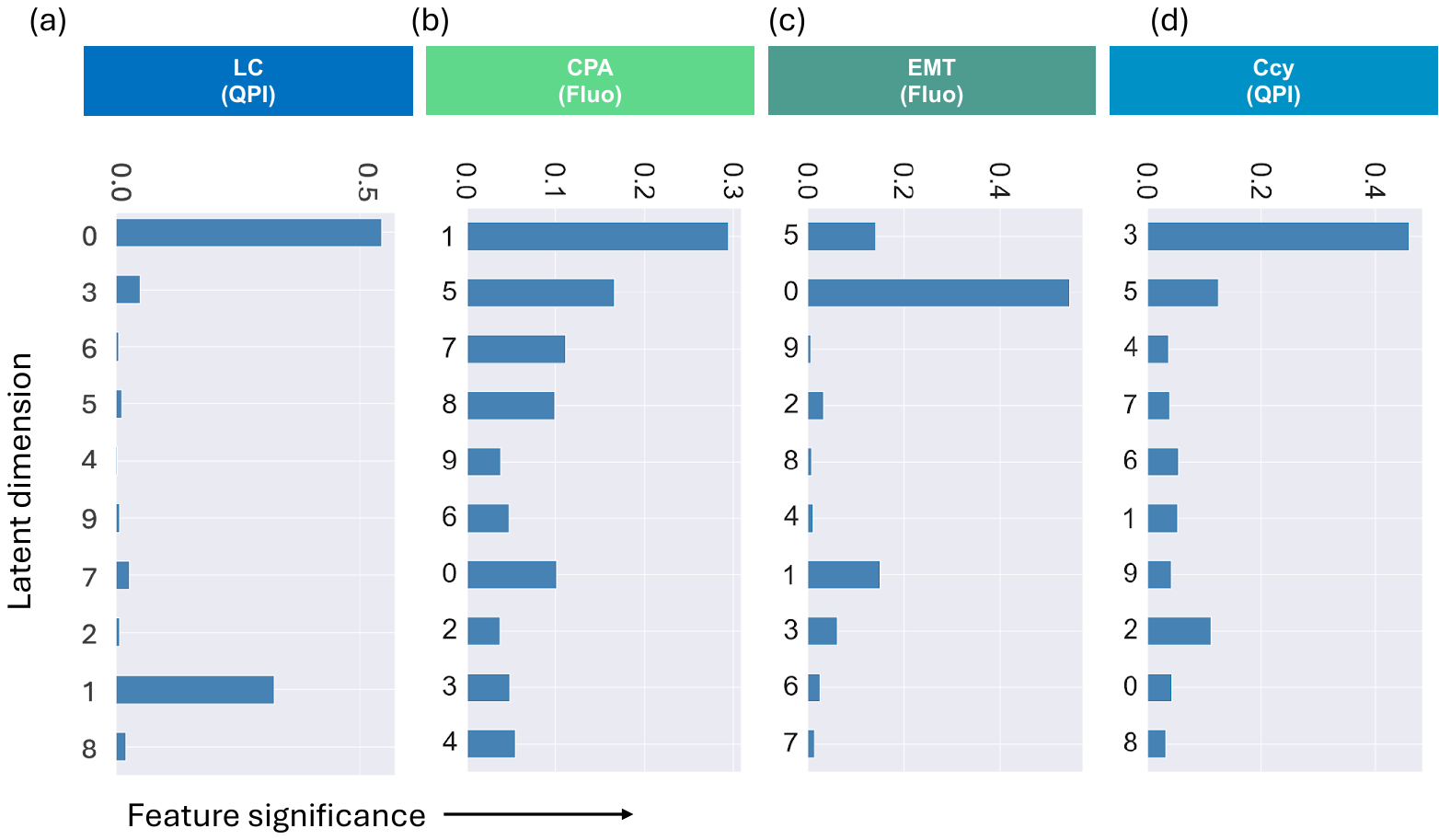

**Figure S5.** Analysis of the significance of the latent features learnt from MorphoGenie model trained with various datasets: LC, Ccy, CPA and EMT.

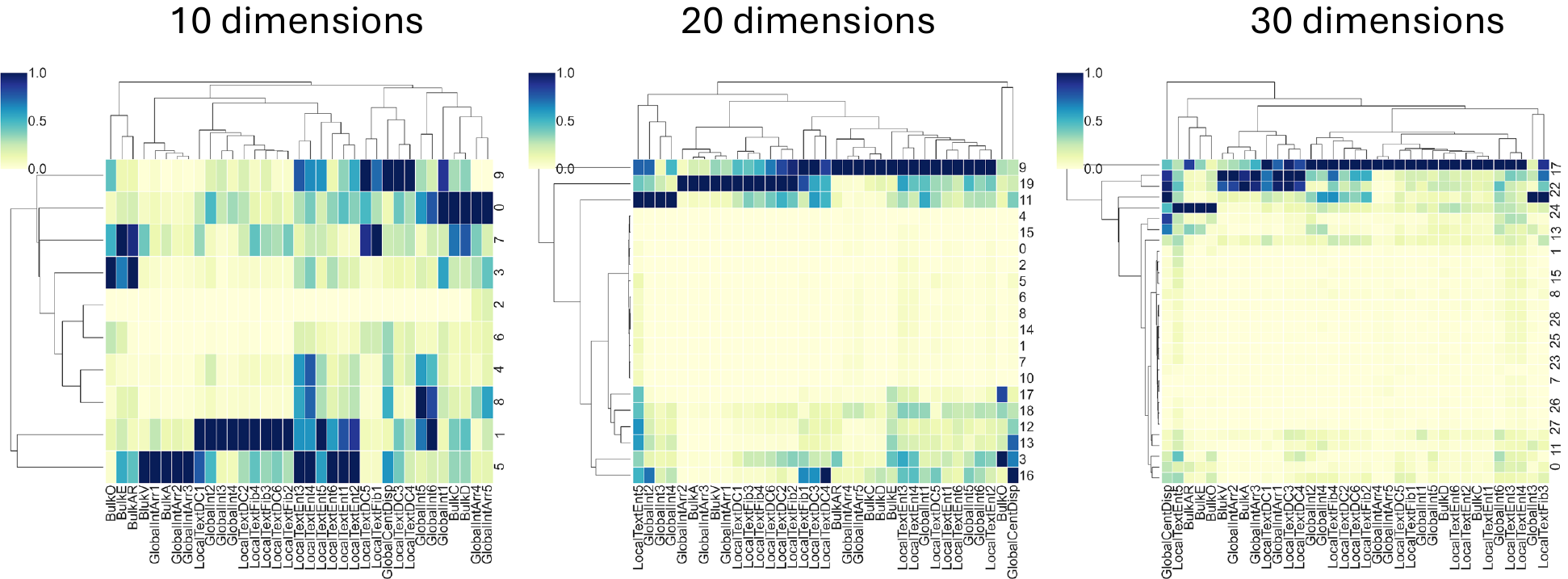

**Figure S6.** Interpretation heatmap illustrating the features learned across the MorphoGenie models with varying latent space dimensions.

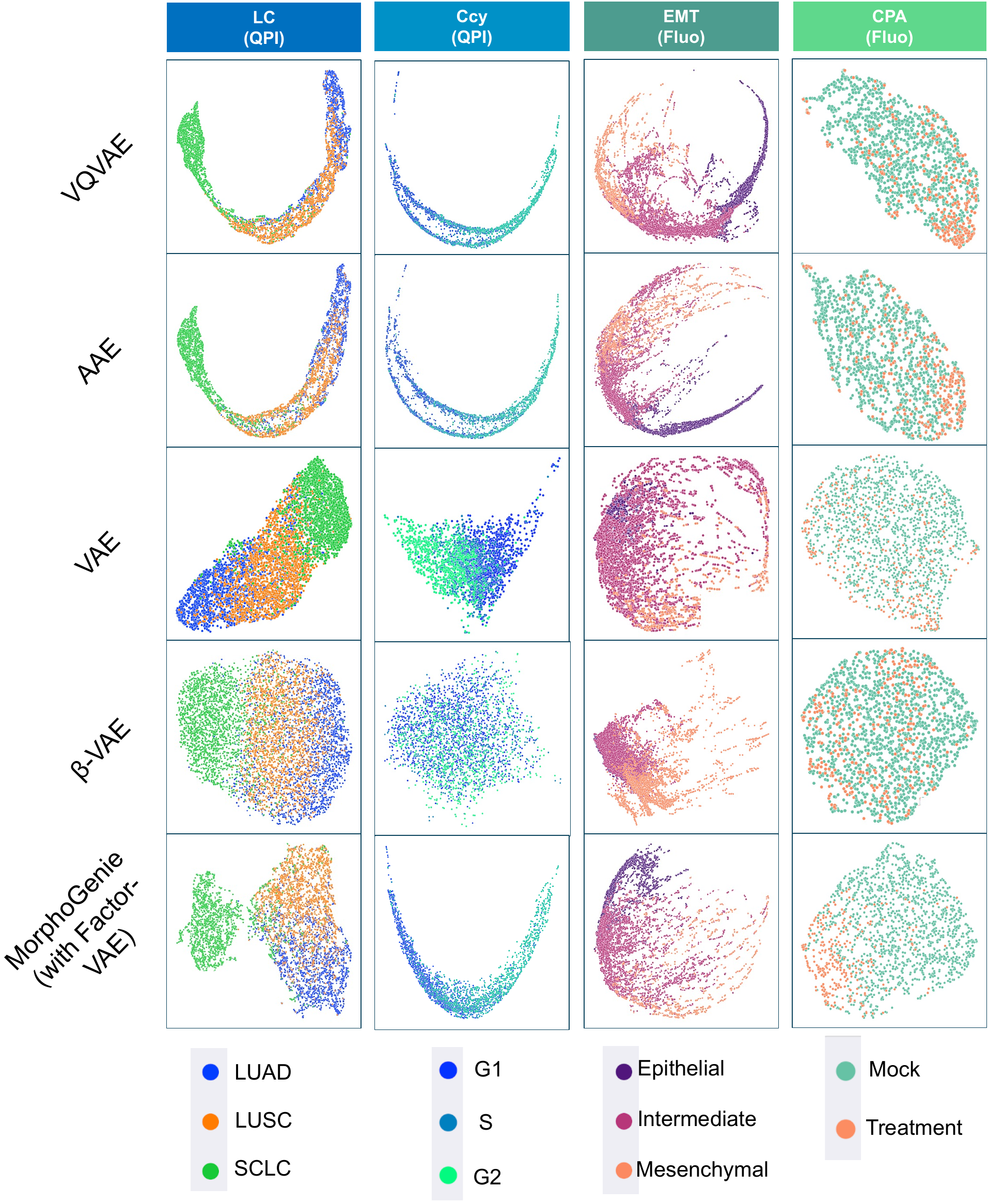

**Figure S7:** Visualization of latent representations across different datasets, based on different models: AQVAE, AAE, VAE, β-VAE and MorphoGenie (with FactorVAE). It demonstrates that MorphoGenie offers advantages of accurate image reconstruction (**Fig. 2, and Supplementary Fig. S2**) without compromising visual data analysis across diverse data types.

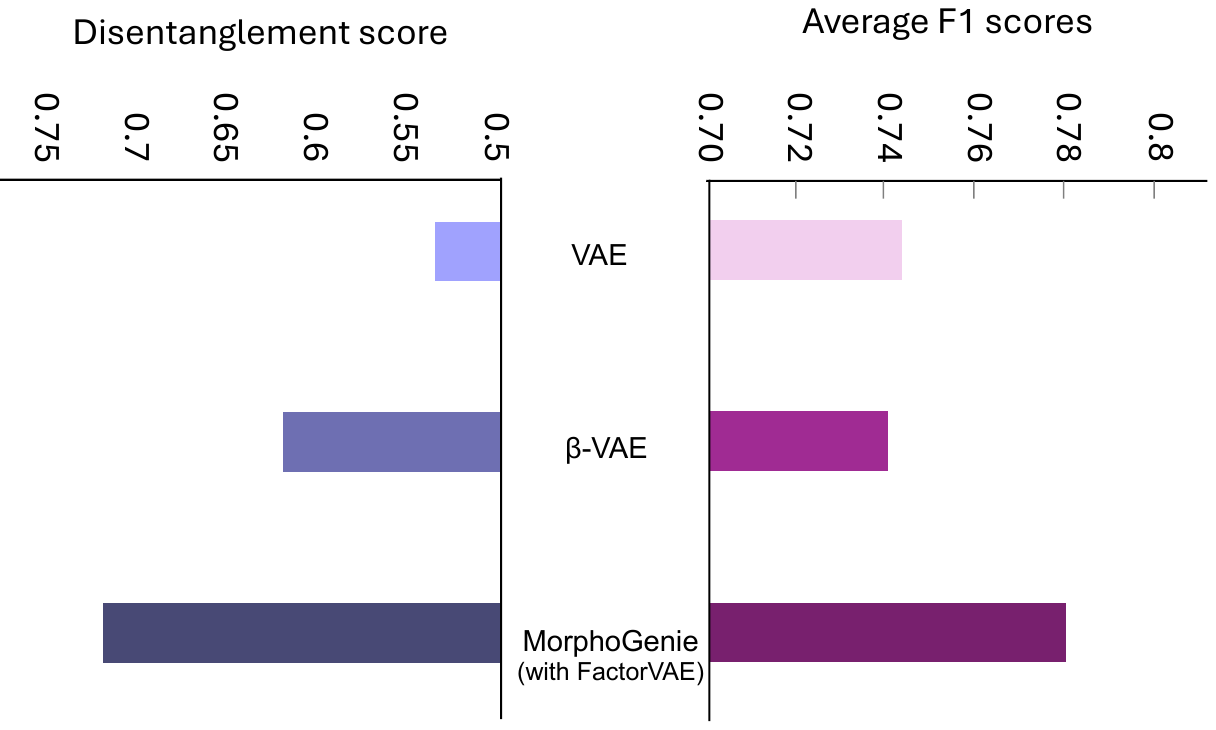

**Figure S8:** Comparisons of the downstream classification performance achieved by different models (based on average F1 scores (right)) across all the datasets included in this study. The disentanglement scores of the models are shown (left) for comparison. It demonstrates that MorphoGenie offers superior classification performance as well as the disentanglement representation learning.

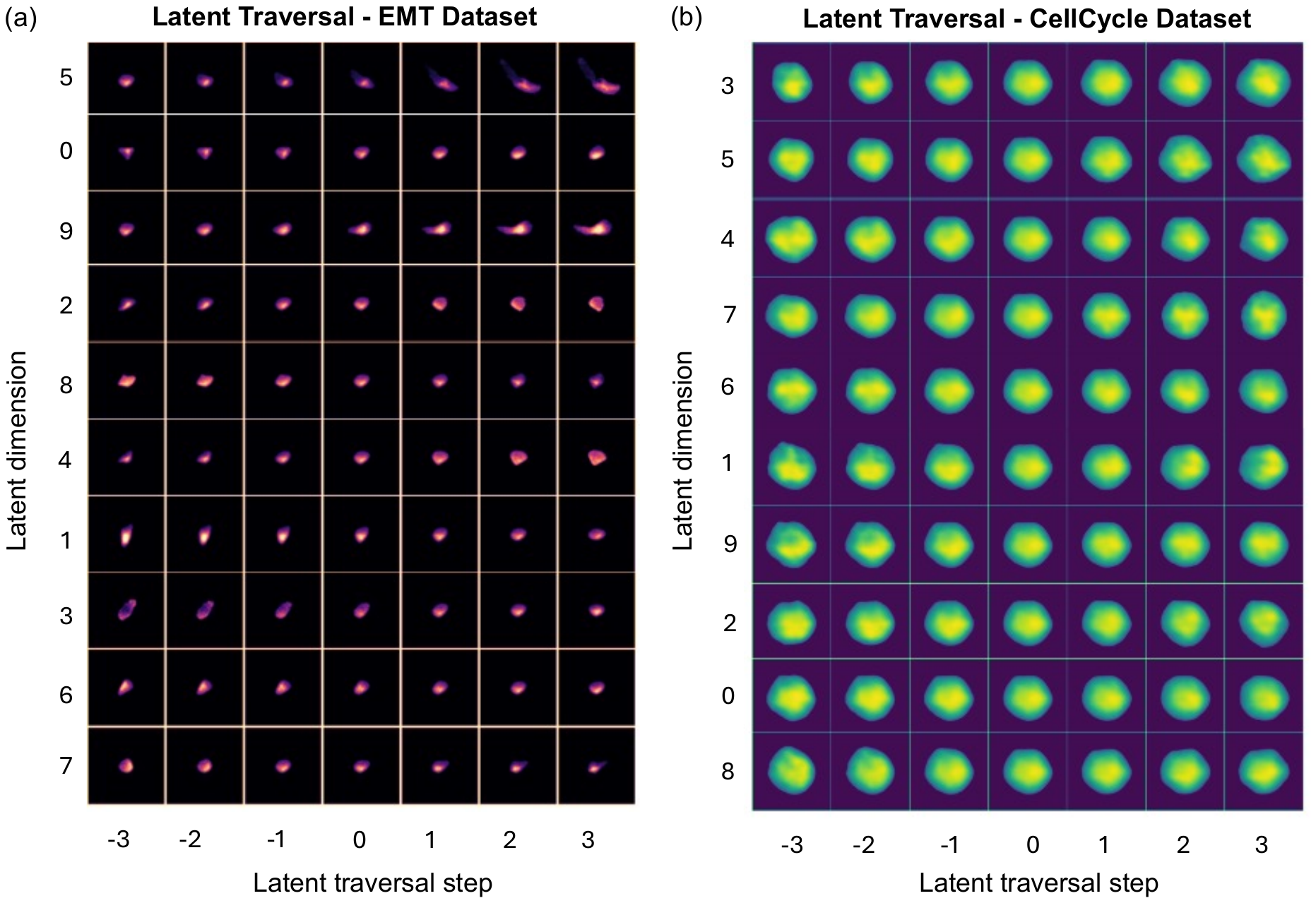

**Figure S9:** Latent traversal reconstructions from the MorphoGenie model trained with (a) the EMT image and (b) cell-cycle datasets. Only 7 traversal steps are shown for clarity. In the interpretation analysis, we actually computed *Q* (=10) steps of latent traversal for each dimension and visualized the morphological variations across each traversal step (See “**Interpretation Heatmap**” in **Methods**).

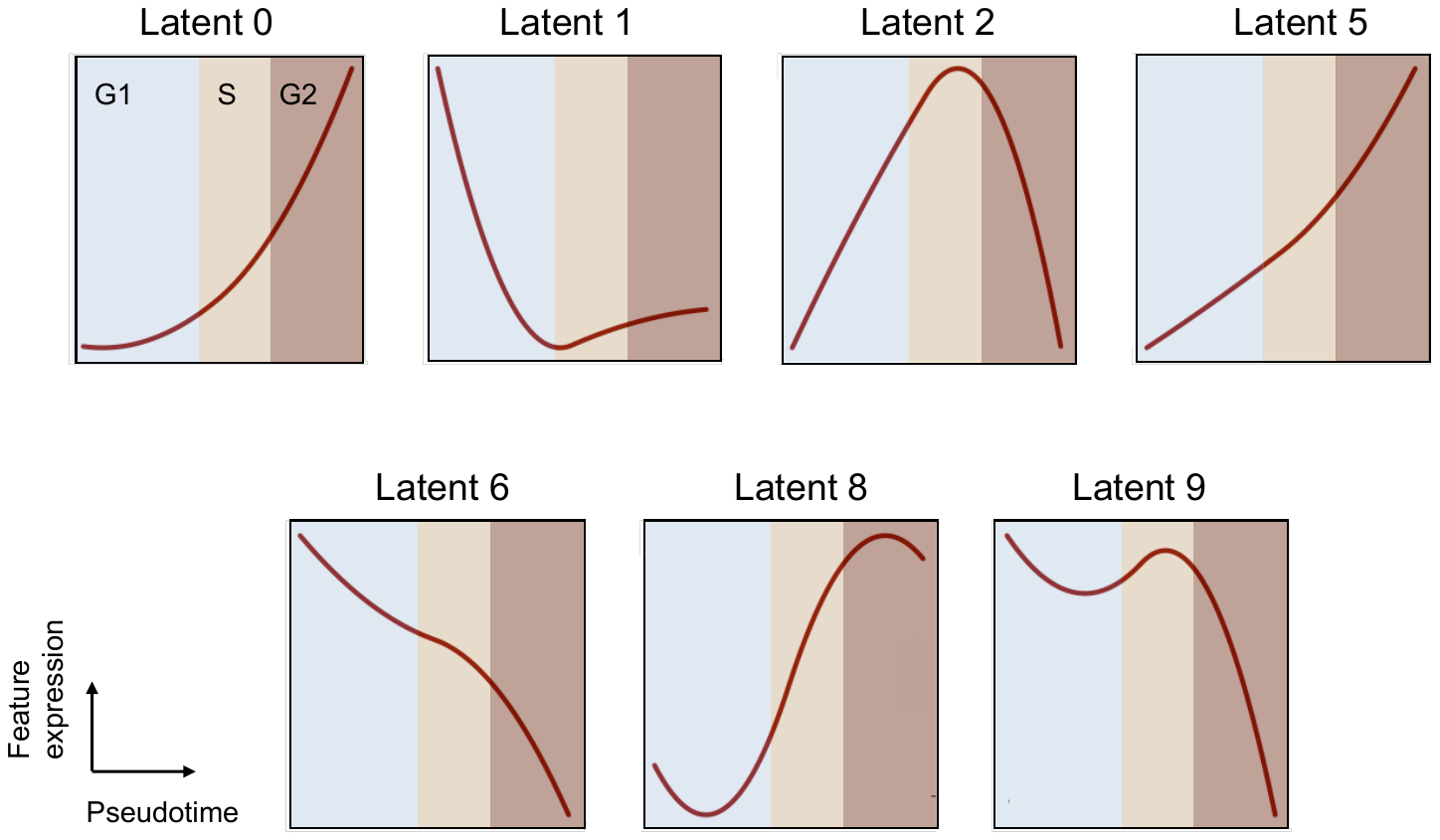

**Figure S10.** Trends of the latent features (based on the FACED cell-cycle imaging dataset) along the pseudotime computed by StaVia.

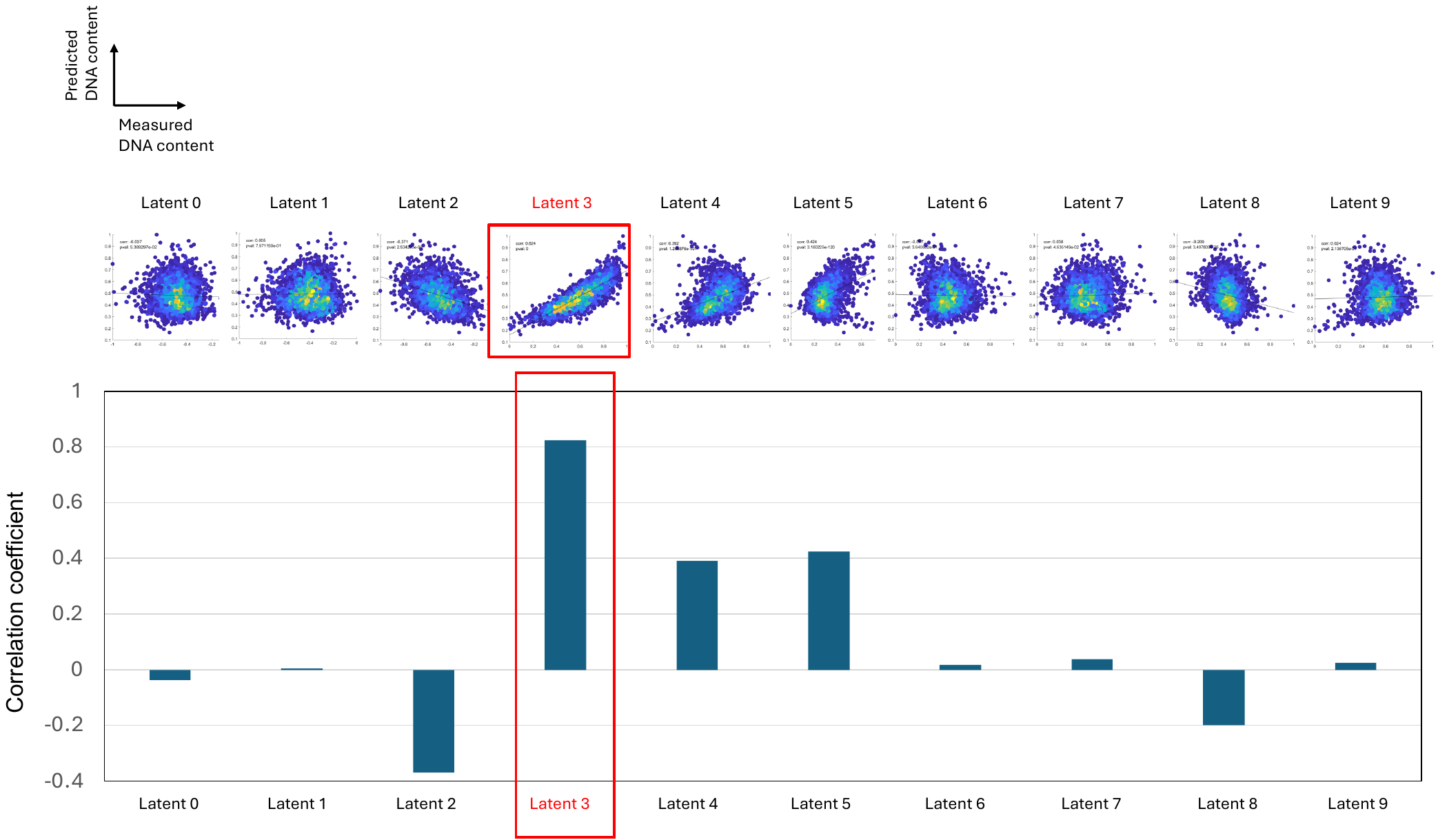

**Figure S11:** Latent-dimension correlative analysis of MorphoGenie's predicted DNA versus the actual DNA measured from the single-cell fluorescence images.

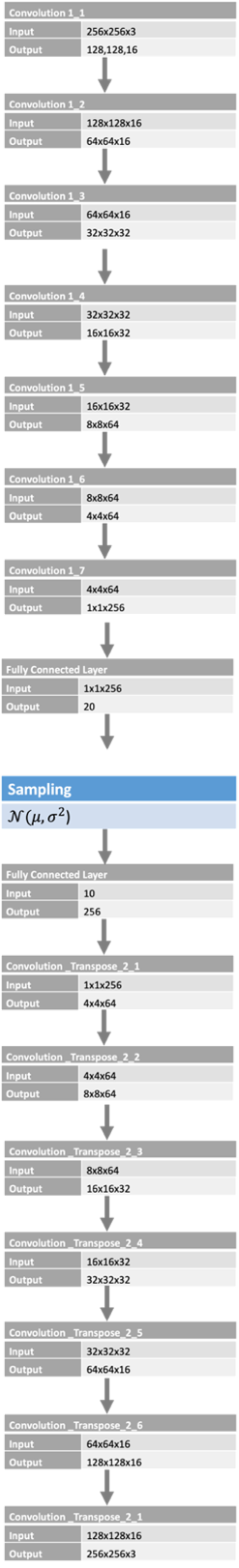

**Figure S12** Architecture of the encoder (before sampling layer) and decoder of FactorVAE (after sampling layer) used in MorphoGenie.

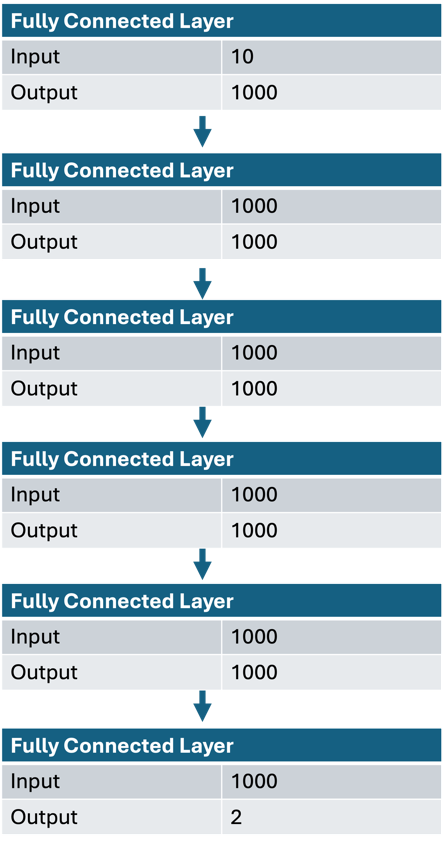

**Figure S13.** Layers in Discriminator of the FactorVAE used in MorphoGenie. This is used to approximate the density ratio present in the KL term (See **Methods**).

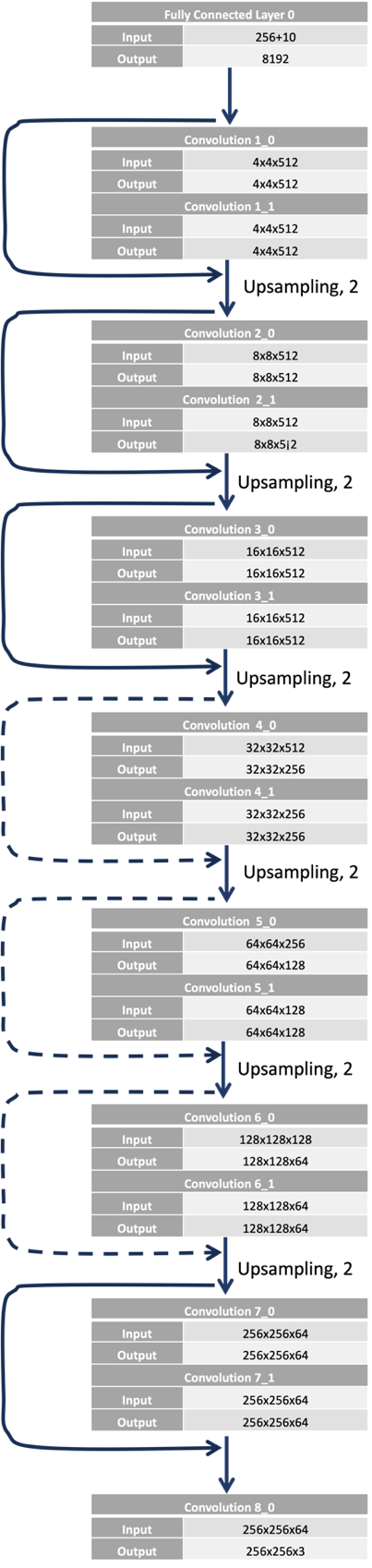

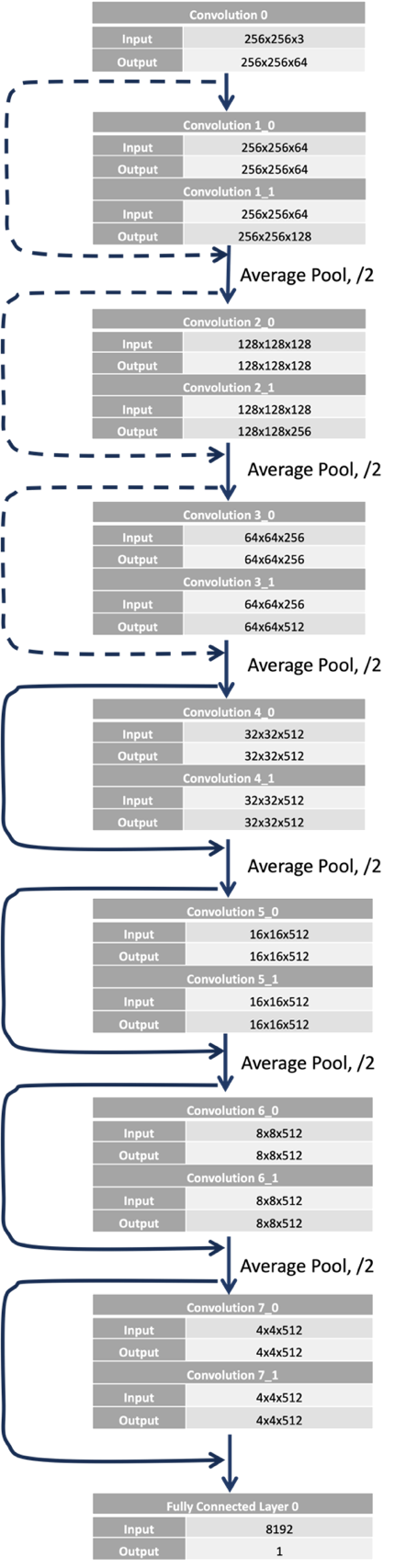

**Figure S14:** Layers of the generator(left) and discriminator (right) of the GAN used in MorphoGenie.

| Bulk Features | | | |
| --- | --- | --- | --- |
| Reference Name | Feature | Symbol | Equation |
| BulkA | Area | A | ${L_{pix}}^{2}\cdot N_{pix}$ |
| BulkV | Volume | V | $\frac{4}{3}\pi\cdot{(\frac{L_{minor}}{2})}^{2}\cdot(\frac{L_{major}}{2})$ |
| BulkC | Circularity |  | $4\pi A/P$ |
| BulkD | Deformation |  |  |
| BulkE | Eccentricity |  | $\frac{L_{ellip}}{L_{major}}$ |
| BulkAR | Aspect Ratio |  | $\frac{L_{minor}}{L_{major}}$ |
| BulkO | Orientation |  | $\theta_{major}$ |

Table S1. List of morphological hierarchical features

| Global Texture Features | | | |
| --- | --- | --- | --- |
| GlobalInt1 | Peak Intensity |  | $max(I)$ |
| GlobalInt2 | Intensity Variance |  | $\frac{\iint_{A} {(I\left( x,y \right)-\bar{\varphi} )}^{2}dxdy}{N_{pix}-1}$ |
| GlobalInt3 | Intensity Skewness |  | $\frac{\iint_{A} {(I\left( x,y \right)-\bar{I} )}^{3}dxdy/N_{pix}}{{\sigma_{Istd}}^{3}}$ |
| GlobalInt4 | Intensity Kurtosis |  | $\frac{\iint_{A} {(I\left( x,y \right)-\bar{I} )}^{4}dxdy/N_{pix}}{{\sigma_{Istd}}^{4}}$ |
| GlobalInt5 | Intensity Range |  | $\max\left\{ I\left( x,y \right) \right\}- min\{I(x,y)\}$ |
| GlobalInt6 | Intensity Minimum |  | $min\{I(x,y)\}$ |
| GlobalRadDist | Intensity Radial Distribution |  | $\frac{\iint_{A} r\cdot I\left( r,\theta\right) drd\theta}{\iint_{A} I\left( r,\theta\right) drd\theta}$ |
| GlobalCentDisp | Intensity Centroid Displacement |  | $\sqrt{{{(x}_{I,cen}-x_{cen})}^{2}+{{(y}_{I,cen}-y_{cen})}^{2}}\cdot L_{pix}$ |
| GlobalIntArr1 | Mean Intensity Arrangement |  | $\frac{\iint_{A} I\left( r,\theta\right) r drd\theta}{\iint_{A} I\left( r,\theta\right) drd\theta}$ |
| GlobalIntArr2 | Intensity Arrangement Variance | ${\sigma_{Iarr}}^{2}$ | $\frac{\iint_{A} {(I\left( r,\theta\right) r)}^{2} drd\theta}{\iint_{A} I\left( r,\theta\right) drd\theta}$ |
| GlobalIntArr3 | Intensity Arrangement Skewness |  | $\frac{\iint_{A} {(I\left( r,\theta\right)\cdot r)}^{3} drd\theta}{{\sigma_{Iarr}}^{2}\cdot\iint_{A} I\left( r,\theta\right) drd\theta}$ |
| GlobalIntArr4 | Intensity Orientation Variance | ${\sigma_{Iang}}^{2}$ | $\frac{\int_{0}^{\infty} {(\tilde{I}(\omega)\cdot\omega)}^{2} d\omega}{\int_{0}^{\infty} \tilde{I}(\omega) d\omega}$ |
| GlobalIntArr5 | Intensity Orientation kurtosis |  | $\frac{\int_{0}^{\infty} {(\tilde{I}(\omega)\cdot\omega)}^{4} d\omega}{{\sigma_{Iang}}^{2}\cdot\int_{0}^{\infty} \tilde{I}(\omega) d\omega}$ |

| Local Texture Features | | | |
| --- | --- | --- | --- |
| LocalTextDC1 | Intensity STD Mean | $\bar{I_{STD}}$ | $\frac{\iint_{A} I_{STD}(x,y) dxdy}{N_{pix}}$ |
| LocalTextDC2 | Intensity STD Variance | ${\sigma_{Istd}}^{2}$ | $\frac{\iint_{A} {(I_{STD}\left( x,y \right)-\bar{I_{STD}} )}^{2}dxdy}{N_{pix}-1}$ |
| LocalTextDC3 | Intensity STD Skewness |  | $\frac{\iint_{A} {(I_{STD}\left( x,y \right)-\bar{I_{STD}} )}^{3}dxdy/N_{pix}}{{\sigma_{Istd}}^{3}}$ |
| LocalTextDC4 | Intensity STD Kurtosis |  | $\frac{\iint_{A} {(I_{STD}\left( x,y \right)-\bar{I_{STD}} )}^{4}dxdy/N_{pix}}{{\sigma_{Istd}}^{4}}$ |
| LocalTextDC5 | Intensity STD Centroid Displacement |  | $\sqrt{{{(x}_{I_{STD},cen}-x_{cen})}^{2}+{{(y}_{I_{STD},cen}-y_{cen})}^{2}}\cdot L_{pix}$ |
| LocalTextDC6 | Intensity STD Radial Distribution |  | $\frac{\iint_{A} r\cdot I_{STD}\left( r,\theta\right) drd\theta}{\iint_{A} I_{STD}\left( r,\theta\right) drd\theta}$ |
| LocalTextEnt1 | Intensity Entropy Mean | $\bar{\varphi_{ent}}$ | $\frac{\iint_{A} I_{ent}(x,y) dxdy}{N_{pix}}$ |
| LocalTextEnt2 | Intensity Entropy Variance | ${\sigma_{\varphi ent}}^{2}$ | $\frac{\iint_{A} {(I_{ent}\left( x,y \right)-\bar{I_{ent}} )}^{2}dxdy}{N_{pix}-1}$ |
| LocalTextEnt3 | Intensity Entropy Skewness |  | $\frac{\iint_{A} {(I_{ent}\left( x,y \right)-\bar{I_{ent}} )}^{3}dxdy/N_{pix}}{{\sigma_{Ient}}^{3}}$ |
| LocalTextEnt4 | Intensity Entropy Kurtosis |  | $\frac{\iint_{A} {(I_{ent}\left( x,y \right)-\bar{I_{ent}} )}^{4}dxdy/N_{pix}}{{\sigma_{Ient}}^{4}}$ |
| LocalTextEnt5 | Intensity Entropy Centroid Displacement |  | $\sqrt{{{(x}_{I_{ent},cen}-x_{cen})}^{2}+{{(y}_{I_{ent},cen}-y_{cen})}^{2}}\cdot L_{pix}$ |
| LocalTextEnt6 | Intensity Entropy Radial Distribution |  | $\frac{\iint_{A} r\cdot I_{ent}\left( r,\theta\right) drd\theta}{\iint_{A} I_{ent}\left( r,\theta\right) drd\theta}$ |
| LocalTextFib1 | Intensity Fiber Centroid Displacement |  | $\sqrt{{{(x}_{Ifiber,cen}-x_{cen})}^{2}+{{(y}_{Ifiber,cen}-y_{cen})}^{2}}\cdot L_{pix}$ |
| LocalTextFib2 | Intensity Fiber Radial Distribution |  | $\frac{\iint_{A} r\cdot I_{fiber}\left( r,\theta\right) drd\theta}{\iint_{A} I_{fiber}\left( r,\theta\right) drd\theta}$ |
| LocalTextFib3 | Intensity Fiber Pixel > Upper Percentile |  | $\frac{Number of pixels in I_{fiber}\left( x,y \right)>75th percentile}{N_{pix}}$ |
| LocalTextFib4 | Intensity Fiber Pixel > Median |  | $\frac{Number of pixels in I_{fiber}\left( x,y \right)>median}{N_{pix}}$ |

Table S2. List of variables used in the feature equations

| Variable | Description | Equation |
| --- | --- | --- |
| $\boldsymbol{C}$ | Contour of binary mask |  |
| $\boldsymbol{CM}$ | Cell mask function | $CM(x,y)=\left\{ \begin{matrix} 1 \\ 0 \end{matrix} \begin{matrix} if inside cell \\ otherwise \end{matrix} \right.$ |
| $\boldsymbol{L}_{\boldsymbol{ellip}}$ | Distance between foci of ellipse |  |
| $\boldsymbol{L}_{\boldsymbol{major}}$ | Major axis length |  |
| $\boldsymbol{L}_{\boldsymbol{minor}}$ | Minor axis length |  |
| $\boldsymbol{L}_{\boldsymbol{pix}}$ | Physical length of one pixel |  |
| $\boldsymbol{I}$ | Pixel Intensity map | $I\left( x,y \right)$ |
| $\boldsymbol{I}\mathbf{(}\boldsymbol{\theta}\mathbf{)}$ | $I$ projected to polar angle |  |
| $\tilde{\boldsymbol{I}}\mathbf{(}\boldsymbol{\omega}\mathbf{)}$ | $I$ in angular frequency domain | $\tilde{I}\left( \omega\right)\mathcal{=F(}I\left( \theta\right))$ |
| $\bar{\boldsymbol{I}_{\boldsymbol{STD}\mathbf{,}\boldsymbol{ker}}}\mathbf{(}\boldsymbol{x}\mathbf{,}\boldsymbol{y}\mathbf{)}$ | Mean value of Intensity within STD filter kernel | $\frac{\int_{x-w_{STD}/2}^{x+w_{STD}/2} \int_{y-w_{STD}/2}^{y+w_{STD}/2} I\left( u,v \right) dvdu}{{w_{STD}}^{2}}$ |
| $\boldsymbol{I}_{\boldsymbol{STD}}\mathbf{(}\boldsymbol{x}\mathbf{,}\boldsymbol{y}\mathbf{)}$ | Intensity STD map | $\int_{x-w_{STD}/2}^{x+w_{STD}/2} \int_{y-w_{STD}/2}^{y+w_{STD}/2} \sqrt{\frac{{(I\left( u,v \right)-\bar{I_{STD,ker}}(x,y))}^{2}}{{w_{STD}}^{2}}} dvdu$ |
| $\boldsymbol{I}_{\boldsymbol{ent}}\mathbf{(}\boldsymbol{x}\mathbf{,}\boldsymbol{y}\mathbf{)}$ | Entropy filtered Intensity | $\sum_{k=0}^{255} p_{I,k}\cdot\log_{2} p_{I,k}$ |
| $\boldsymbol{I}_{\boldsymbol{fiber}}\mathbf{(}\boldsymbol{x}\mathbf{,}\boldsymbol{y}\mathbf{)}$ | Fiber texture enhanced Intensity | $FF(I\left( x,y \right))$ |
| $\boldsymbol{N}_{\boldsymbol{pix}}$ | Pixel number in cell mask | $\iint CM(x,y) dA$ |
| $\boldsymbol{P}$ | Perimeter | $\oint_{C} \sqrt{\left( \left( \frac{dx}{d\theta} \right)^{2}+\left( \frac{dy}{d\theta} \right)^{2} \right)}d\theta$ |
| $\boldsymbol{p}_{\boldsymbol{I}\mathbf{,}\boldsymbol{k}}\mathbf{(}\boldsymbol{x}\mathbf{,}\boldsymbol{y}\mathbf{)}$ | Normalized histogram counts within kernel of $I$ | $p_{I,k}\left( x,y \right)=\frac{number of pixels in kernel \left( w_{ent} \right) with I=k}{Total number of pixels in kernel}$ |
| $\boldsymbol{r}\mathbf{,}\boldsymbol{\theta}$ | Polar coordinates centered at cell centroid |  |
| $\boldsymbol{w}_{\boldsymbol{ent}}$ | Kernel size of entropy filter |  |
| $\boldsymbol{w}_{\boldsymbol{STD}}$ | Kernel size of STD filter |  |
| $\boldsymbol{x}\mathbf{,}\boldsymbol{y}$ | Cartesian coordinates |  |
| $\boldsymbol{x}_{\boldsymbol{cen}}$  $\boldsymbol{y}_{\boldsymbol{cen}}$ | Coordinates of cell centroid | $x_{cen}=\frac{\iint_{A} x\cdot CM\left( x,y \right) dxdy}{N_{pix}}$  $y_{cen}=\frac{\iint_{A} y\cdot CM\left( x,y \right) dxdy}{N_{pix}}$ |
| $\boldsymbol{x}_{\boldsymbol{I}\mathbf{,}\boldsymbol{cen}}$  $\boldsymbol{y}_{\boldsymbol{I}\mathbf{,}\boldsymbol{cen}}$ | Coordinates of $I$ weighted cell centroid | $x_{MD,cen}=\frac{\iint_{A} x\cdot I\left( x,y \right) dxdy}{N_{pix}}$  $y_{MD,cen}=\frac{\iint_{A} y\cdot I\left( x,y \right) dxdy}{N_{pix}}$ |
| $\boldsymbol{x}_{\boldsymbol{Ient}\mathbf{,}\boldsymbol{cen}}$  $\boldsymbol{y}_{\boldsymbol{Ient}\mathbf{,}\boldsymbol{cen}}$ | Coordinates of entropy filtered $I$ weighted cell centroid | $x_{Ient,cen}=\frac{\iint_{A} x\cdot I_{ent}\left( x,y \right) dxdy}{N_{pix}}$  $y_{Ient,cen}=\frac{\iint_{A} y\cdot I_{ent}\left( x,y \right) dxdy}{N_{pix}}$ |
| $\boldsymbol{x}_{\boldsymbol{Ifiber}\mathbf{,}\boldsymbol{cen}}$  $\boldsymbol{y}_{\boldsymbol{Ifiber}\mathbf{,}\boldsymbol{cen}}$ | Coordinates of fiber enhanced $I$ weighted cell centroid | $x_{Ifiber,cen}=\frac{\iint_{A} x\cdot I_{fiber}\left( x,y \right) dxdy}{N_{pix}}$  $y_{Ifiber,cen}=\frac{\iint_{A} y\cdot I_{fiber}\left( x,y \right) dxdy}{N_{pix}}$ |
| $\boldsymbol{x}_{\boldsymbol{ISTD}\mathbf{,}\boldsymbol{cen}}$  $\boldsymbol{y}_{\boldsymbol{ISTD}\mathbf{,}\boldsymbol{cen}}$ | Coordinates of STD filtered $I$ weighted cell centroid | $x_{ISTD,cen}=\frac{\iint_{A} x\cdot I_{STD}\left( x,y \right) dxdy}{N_{pix}}$  $y_{ISTD,cen}=\frac{\iint_{A} y\cdot I\left( x,y \right) dxdy}{N_{pix}}$ |
